# The breast cancer oncogene IKKε coordinates mitochondrial function and serine metabolism

**DOI:** 10.1101/855361

**Authors:** Ruoyan Xu, William Jones, Ewa Wilcz-Villega, A. Sofia H. Costa, Vinothini Rajeeve, Robert B. Bentham, Kevin Bryson, Ai Nagano, Busra Yaman, Sheila Olendo Barasa, Yewei Wang, Claude Chelala, Pedro Cutillas, Gyorgy Szabadkai, Christian Frezza, Katiuscia Bianchi

## Abstract

The IκB kinase ε (IKKε) is a key molecule at the crossroads of inflammation and cancer. Known for its role as an activator of NFκB and IRF3 signalling leading to cytokine secretion, the kinase is also a breast cancer oncogene, overexpressed in a variety of tumours. However, to what extent IKKε remodels cellular metabolism is currently unknown. Here we used a combination of metabolomics and phosphoproteomics to show that IKKε orchestrates a complex metabolic reprogramming that affects mitochondrial metabolism and serine biosynthesis. Acting independently of its canonical signalling role, IKKε upregulates the serine biosynthesis pathway (SBP) mainly by limiting glucose and pyruvate derived anaplerosis of the TCA cycle. In turn, this elicits activation of the transcription factor ATF4 and upregulation of the SBP genes. Importantly, pharmacological inhibition of the IKKε-induced metabolic phenotype reduces proliferation of breast cancer cells. Finally, we show that in a set of basal ER negative and highly proliferative human breast cancer tumours, IKKε and PSAT1 expression levels are positively correlated corroborating the link between IKKε and the SBP in the clinical context.

## INTRODUCTION

Chronic inflammation, triggered by the tumour stroma or driven by oncogenes, plays a central role in tumour pathogenesis (Netea *et al*, 2017). A key step leading to inflammation in both compartments is activation of the transcription factor Nuclear Factor κB (NFκB), mediated via canonical or alternative, non-canonical pathways. Key players in both pathways are the members of the IκB kinase (IKK) family, which, by phosphorylating IκB, induce its proteasome-mediated degradation, a step required for releasing NFκB from the IκB-imposed cytosolic localization and thus leading to its nuclear translocation (Clément *et al*, 2008).

Evidence in support of the crucial role played by the IKK family in inflammation-induced malignant transformation was provided by the reduction of tumour incidence following the deletion of the canonical IKK family member IKKβ in intestinal epithelial and myeloid cells in a mouse model of colitis associated cancer development (Greten *et al*, 2004). Soon after, the non-canonical member of the IKK family, IKKε, was shown to induce breast cancer (Boehm *et al*, 2007) and to be overexpressed in ovarian (Guo *et al*, 2009), prostate (Péant *et al*, 2011) and non-small cell lung cancers (Guo *et al*, 2013), pancreatic ductal carcinoma (Cheng *et al*, 2011), and glioma (Guan *et al*, 2011). In particular, IKKε was shown to induce breast cancer via mechanisms involving CYLD (Hutti *et al*, 2009) and TRAF2 (Zhou *et al*, 2013), ultimately mediating NFκB activation (Boehm *et al*, 2007).

Beyond cancer, IKKε is a key regulator of innate and adaptive immunity, activating NFκB and Interferon Regulatory factor 3 (IRF3), inducing type I interferon (Clément *et al*, 2008; Zhang *et al*, 2016), but activation of the interferon response has been reported not to be essential for IKKε-mediated cellular transformation (Boehm *et al*, 2007). On the other hand, IKKε has been shown to regulate central carbon metabolism both in immune and cancer cells. In dendritic cells (DCs), IKKε, together with its closest homologue TANK binding kinase 1 (TBK1), is required for the switch to aerobic glycolysis induced by activation of the Toll-like receptors (TLRs) and activation of DC. Glycolysis is the main glucose catabolic pathway, whereby through a series of reactions cells metabolise glucose to pyruvate, which, in the presence of oxygen, is in turn oxidised to CO_2_ in the mitochondrial matrix via the TCA cycle to produce ATP using the mitochondrial respiratory chain. Lack of oxygen prevents the mitochondrial utilization of pyruvate, and glucose is converted into lactate (anaerobic glycolysis). In contrast, aerobic glycolysis refers to a metabolic condition whereby glucose is not fully oxidised in the mitochondria, even in the presence of oxygen, and is utilised for the production of amino acids, lipids and nucleotides via pathways branching out from the glycolysis and TCA cycle. Accordingly, aerobic glycolysis in DCs allows fatty acids synthesis, which is required for the expansion of the endoplasmic reticulum and Golgi, supporting DC activation (Everts *et al*, 2014). Allowing the production of key cellular constituents, aerobic glycolysis is most frequently observed in highly proliferative cells, such as activated immune cells and cancer cells (Andrejeva & Rathmell, 2017). Accordingly, IKKε also regulates glucose uptake in pancreatic ductal adenocarcinoma as well as mitochondrial function in mouse embryonic fibroblasts (MEFs) (Zubair *et al*, 2016; Reilly *et al*, 2013). However, a comprehensive investigation of the role of IKKε as regulator of cellular metabolism in cancer has not yet been carried out.

The serine biosynthesis pathway (SBP) is known to be dysregulated in cancer as target of a series of oncogenes (Amelio *et al*, 2014; Yang & Vousden, 2016), and phosphoglycerate dehydrogenase (PHGDH), the first enzyme of the pathway, is amplified in breast cancer and melanoma, where it functions as an oncogene (Locasale *et al*, 2011; Possemato *et al*, 2011). However, whether inflammatory signalling regulates the SBP pathway to promote tumorigenesis has not yet been investigated.

Considering the emerging role of IKKε in inflammation and breast tumorigenesis, where the SBP has a known pro-oncogenic role, we investigated the potential intersection between IKKε−mediated inflammatory pathways and the SBP. We found that IKKε directly controls the SBP via activation of a mitochondria-nuclear retrograde pathway and the transcription factor ATF4, leading to upregulation of the SBP enzymes, in particular phosphoserine aminotransferase 1 (PSAT1). Importantly we also demonstrate that IKKε-mediated regulation of cellular metabolism is independent of the canonical signalling pathway via NFκB/IRF3. Moreover, we have identified a subset of basal, estrogen receptor negative (ER^-^) highly proliferative breast tumours where IKKε and PSAT1 expression correlates, confirming the pathophysiological role of our findings. These results identify an additional role for IKKε in breast cancer, adding regulation of cellular metabolism to the canonical oncogenic pathways. Thus, our data suggest a synergistic mechanism of action by which alterations of cellular metabolism and inflammation driven by the IKKε oncogene support tumour growth and proliferation.

## RESULTS

### IKKε rewires cellular metabolism

To investigate the effect of IKKε activation on metabolism, we used two cellular model systems: (i) doxycycline inducible Flp-In 293 HA-IKKε-expressing cells and their respective GFP expressing controls (Flp-In 293 HA-GFP cells) and (ii) two breast cancer cell lines, T47D and MDA-MB-468, where the kinase was silenced via siRNA. HEK-293 cells do not express endogenous IKKε, thus we could modulate its expression to a level that matched those observed in breast cancer cell lines, where IKKε is highly expressed (Fig. 1A and ref. (Boehm *et al*, 2007)). Liquid chromatography-mass spectrometry (LC-MS) analysis of steady state metabolite levels revealed that induction of IKKε expression affected a large fraction of the measured metabolites (26 out of 32, Fig. 1B and Table 1). To account for any possible effect of doxycycline on cell metabolism, we compared cells with doxycycline-induced expression of IKKε versus GFP (Ahler *et al*, 2013). Of particular interest, IKKε increased cellular glucose and glutamine levels, along with a group of amino acids, including serine and glycine. Conversely, IKKε reduced steady state levels of TCA cycle intermediates (e.g. citrate and malate). The increased intracellular level of serine was a consequence of increased biosynthesis as we observed a significant increase in the level of ^13^C_6_-glucose-derived serine (m+3 isotopologue), suggesting that IKKε positively regulates the SBP (Fig. 1C). A key enzyme in the SBP is phosphoserine amino-transferase 1 (PSAT1), which transfers nitrogen from glutamine-derived glutamate to phosphohydroxypyruvate, generating phosphoserine for the final dephosphorylation step of serine biosynthesis (Fig. 1D). Using ^15^N_2_-glutamine labelling, we confirmed increased levels of nitrogen labelling of serine (m+1) in IKKε expressing cells (see Fig. 1C and Table 1), consistent with an increase in PSAT1 transamination activity, corroborating that serine biosynthesis was activated by IKKε. In contrast, we observed a significant reduction of citrate m+2 and malate m+2 accumulation from ^13^C_6_-glucose, indicating that IKKε reduces the activity of the pyruvate dehydrogenase (PDH) (Fig. 1E). Fractional enrichment analysis for these metabolites showed that together with an increased serine biosynthesis, IKKε expression increased the uptake of unlabelled serine from the media, causing also an increase in the m+0 isotopologue (Fig. 1F). Moreover, the fraction of ^13^C_6_-glucose derived citrate and malate (m+2, pyruvate dehydrogenase generated) was reduced in IKKε expressing cells, causing reduction in their total levels, indicating that no other carbon sources (e.g. glutamine) compensate for the lack of pyruvate entering the TCA cycle (Fig. 1F). The fraction of ^15^N labelled serine, derived from glutamine was also increased, indicating higher glutamine usage in serine biosynthesis as nitrogen source (Fig. 1G).

**Figure 1.**
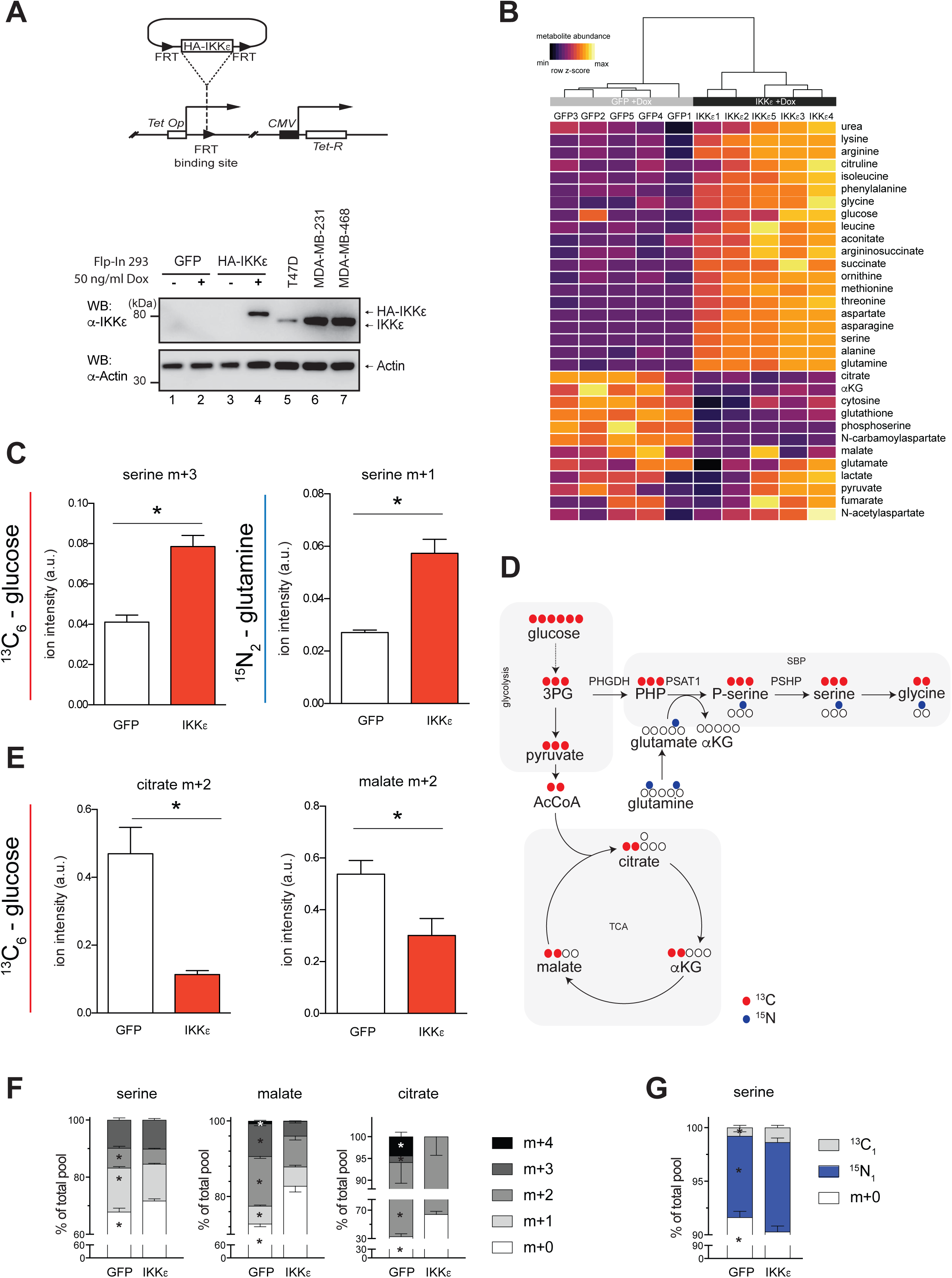
IKKε induces remodelling of cellular carbon metabolism by activating the serine biosynthesis pathway (SBP) and suppressing pyruvate oxidation. (A) Top panel: Scheme illustrating the tetracycline inducible Flp-In 293 system that controls the expression of HA-IKKε or HA-GFP. Bottom panel: Induced expression of HA-IKKε in Flp-In 293 cells treated with doxycycline for 16 hours compared to endogenous IKKε in T47D, MDA-MB-231 and MDA-MB-468 breast cancer cell lines. (B) Heatmap and hierarchical clustering of metabolite concentrations in Flp-In 293 HA-GFP and Flp-In 293 HA-IKKε cells treated with doxycycline (50 ng/ml, 16 hours) (n=5) (C) Serine production from glucose (serine m+3, ^13^C_6_-glucose labelling, left panel) and glutamine (serine m+1, ^15^N_2_-glutamine labelling, right panel) in Flp-In 293 HA-GFP or Flp-In 293 HA-IKKε cells treated with doxycycline (50 ng/ml, 16 hours). (n=5) (D) Schematic representation of the ^13^C_6_-glucose and ^15^N_2_-glutamine labelling strategy to assess the effect of HA-IKKε induction on glycolysis, the TCA cycle and serine metabolism. (E) Contribution of pyruvate and glucose-derived carbon to TCA cyle metabolites in Flp-In 293 HA-GFP or Flp-In 293 HA-IKKε cells treated with doxycycline (50 ng/ml, 16 hours). (n=5). (F) Fractional enrichment of serine, malate and citrate ^13^C-isotopologues in Flp-In 293 HA-GFP and Flp-In 293 HA-IKKε cells treated with doxycycline (50 ng/ml, 16 hours). (G) Fractional enrichment of the serine ^15^-N-isotopologue in Flp-In 293 HA-GFP and Flp-In 293 HA-IKKε cells treated with doxycycline (50 ng/ml, 16 hours). m+1 shows the naturally occurring ^13^C isotopologue. Data Information: In (C, E-G), metabolite levels were normalised to the internal standard HEPES. In (C) and (E), data are presented as mean ± SD. In (C) and (E-G), *p<0.05 (two-tailed Student’s t-test).

We then investigated whether IKKε has a similar metabolic function in breast cancer cell lines, where it is constitutively expressed. Because IKKε has been shown to be an oncogene across different breast cancer subtypes (Boehm *et al*, 2007), we decided to use the T47D and the MDA-MB-468 cell lines, modelling estrogen receptor positive (ER^+^) and triple negative breast cancer, respectively (Xing *et al*). After successful silencing of the kinase (Fig. 2A,B), ^13^C_6_-glucose and ^15^N_2_-glutamine labelling analysis confirmed the overall effect of IKKε on cellular metabolism. In particular on the serine and glycine biosynthesis pathways, IKKε silencing exerted the opposite effect to HA-IKKε induction in the Flp-In 293 model (Fig. 2C-F and Table 1). Along these lines, IKKε knock-down resulted in increased levels of the TCA cycle metabolites citrate and malate m+2 isotopologues, derived from ^13^C-glucose via PDH (Fig. 2E,F). Taken together, these data indicate that in cancer cells IKKε redirects a significant fraction of glucose-derived carbons to the SBP and reduces pyruvate oxidation in the TCA cycle.

**Figure 2.**
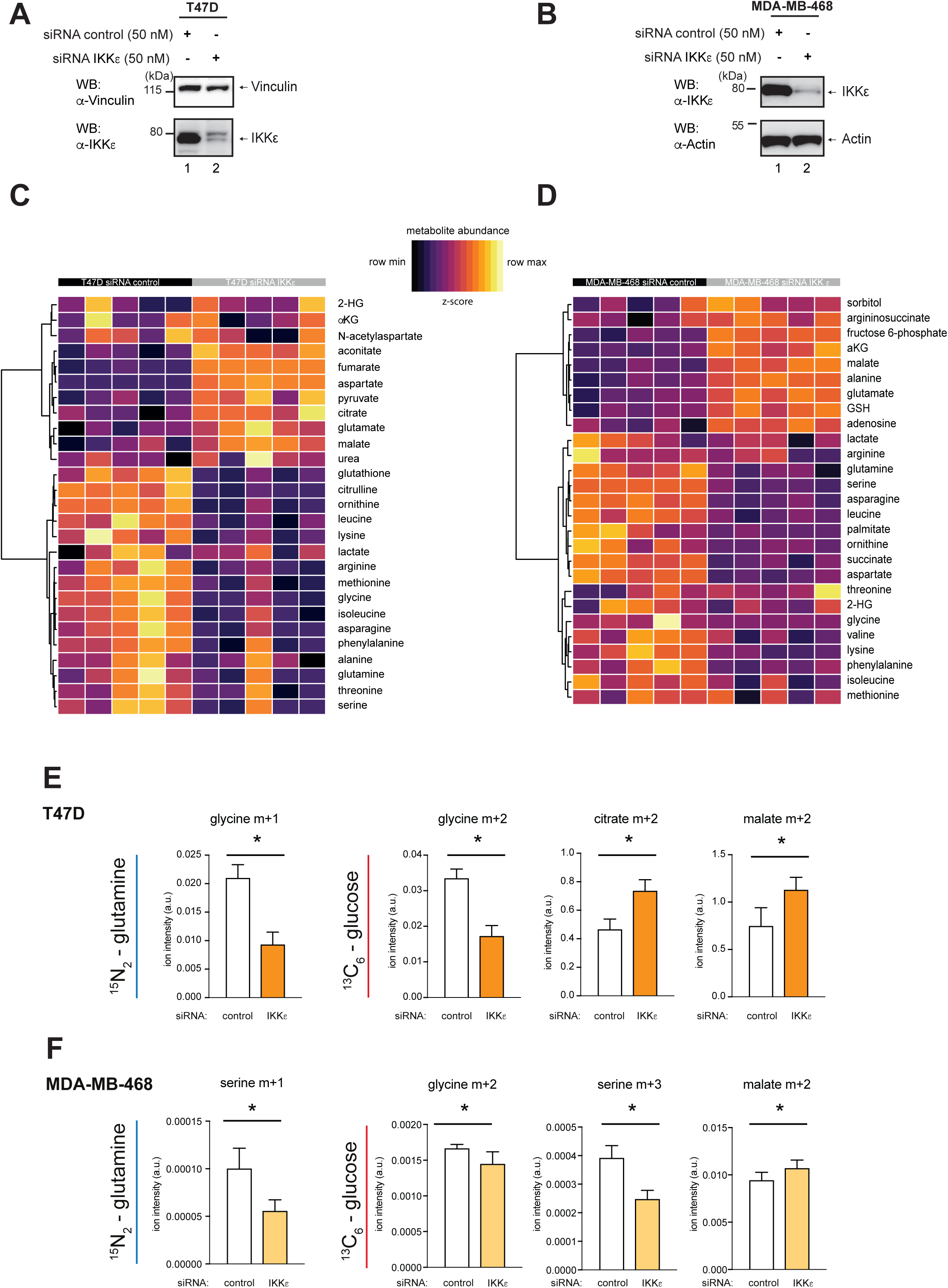
The effect of IKKε silencing on metabolism of breast cancer cell lines. (A-B) Level of IKKε in *IKBKE*-silenced (A) T47D and (B) MDA-MB-468 breast cancer cell lines. (C-D) Heatmap and hierarchical clustering of metabolite concentrations in (C) *IKBKE*-silenced T47D cells and (D) *IKBKE-*silenced MDA-MB-468 cells. (E) Glycine production, representative of serine production, from glutamine (glycine m+1, ^15^N-glutamine labelling) and glucose (glycine m+2, ^13^C_6_-glucose labelling), and contribution of pyruvate and glucose-derived carbon to TCA cycle metabolites (citrate m+2, malate m+2, ^13^C_6_-glucose labelling) in *IKBKE*-silenced T47D cells. (n=5). (F) Serine production from glutamine (serine m+1, ^15^N-glutamine labelling), and serine and glycine production from glucose (glycine m+2, serine m+3, ^13^C_6_-glucose labelling) as well as contribution of pyruvate and glucose-derived carbon to TCA cycle metabolites (malate m+2, ^13^C_6_-glucose labelling) in *IKBKE*-silenced MDA-MB-468 cells. (n=5). Data Information: In (C, E), metabolite levels were normalised to the internal standard HEPES. In (D, F), metabolite levels were normalised to total ion count. In (C-D) metabolite levels were scaled to maximum and minimum levels of each metabolite. In (E-F), *p<0.05 (two-tailed Student’s t-test)

### IKKε contributes to PSAT1 phosphorylation

IKKε is a Ser/Thr protein kinase. Therefore, in order to understand how it regulates cellular metabolism, we compared the phosphoproteomes of three independent control (GFP) and IKKε expressing Flp-In 293 clones. Multivariate analysis showed that the two clones highly expressing IKKε grouped together in principal component analysis (PCA) and were separated from controls and cells expressing IKKε at low levels (Fig. 3A, B). These results suggested that IKKε induces a dose-dependent effect in the phosphoproteome of these cells. Pathway enrichment analysis indicated that in addition to the known innate immunity pathways (KEGG: hepatitis B/hsa05161 and Epstein-Barr virus infection/hsa05169), several metabolic pathways were enriched among the differentially phosphorylated substrates (KEGG: biosynthesis of amino acids/hsa01230 and glycolysis/gluconeogenesis/hsa00010) (Fig. 3C). Of the >3000 phosphopeptides quantified in four technical replicates, multiple IKKε residues and PSAT1 at Ser331 were among the most induced phosphorylation sites correlating with IKKε expression (Table 2 and Fig. 3D). Moreover, we also identified mitochondrial substrates, including the E1 subunit of the pyruvate dehydrogenase complex (PDHA1) (Table2). Of note, a PSAT1 phosphorylation motif was highly homologous to a canonical IKKε target motif (Supplementary Fig. S1A,B) (Hutti *et al*, 2009). Therefore, we tested whether PSAT1 was a direct IKKε substrate. We expressed a HA-tagged version of IKKε and extracted it using immunoprecipitation (IP). Proteomic analysis of HA-IKKε pull-downs showed that apart from IKKε, only TANK binding kinase 1 (TBK1), the closest homologue and heterodimer binding partner of IKKε (Chau *et al*, 2008) was binding specifically (Supplementary Fig. S1C), corroborating the validity of the approach. Notably, the identified consensus sequences of TBK1 and IKKε are identical (Hutti *et al,* 2012), and using an *in vitro* kinase assay we showed that a kinase complex, containing IKKε and TBK1, was capable of phosphorylating PSAT1 (S331) (Supplementary Fig. S1D). Interestingly, whilst PSAT1 phosphorylation was also detected in samples containing IKKε kinase-dead mutant (KM - K38A (Ikeda *et al*, 2007)) despite the lack of autophosphorylation of the mutant (Supplementary Fig. S1E), TBK1 alone was not sufficient to induce this effect, confirming that the role of IKKε in PSAT1 phosphorylation is not redundant. The pS331 phospho-site of PSAT1 was previously reported to be phosphorylated in a range of conditions (Klammer *et al*, 2012; Mertins *et al*, 2013, 2014, 2016; Sharma *et al*, 2014; Stuart *et al*, 2015), but to date, no kinase has been identified to mediate this effect. In order to validate PSAT1 phosphorylation in human breast cancer samples, we re-analysed a set of publicly available phoshoproteomic data obtained from patients (Mertins *et al*, 2016). Out of the 77 cases analysed, altered PSAT1 phosphorylation could be observed in 14. Among these, in 8 cases the level was observed as negative (lower than the average), while in 6 cases PSAT1 was phosphorylated. Interestingly, we noticed that all the 8 negative cases were classified as Luminal A or B and were ER/PR positive, while 5 cases where PSAT1 was phosphorylated were classified as Basal and were triple negative tumours. These data indicate that PSAT1 phosphorylation occurs in human breast cancer, and its phosphorylation might be associated with cancer subtypes with poorer clinical outcomes (Table 3).

**Figure 3.**
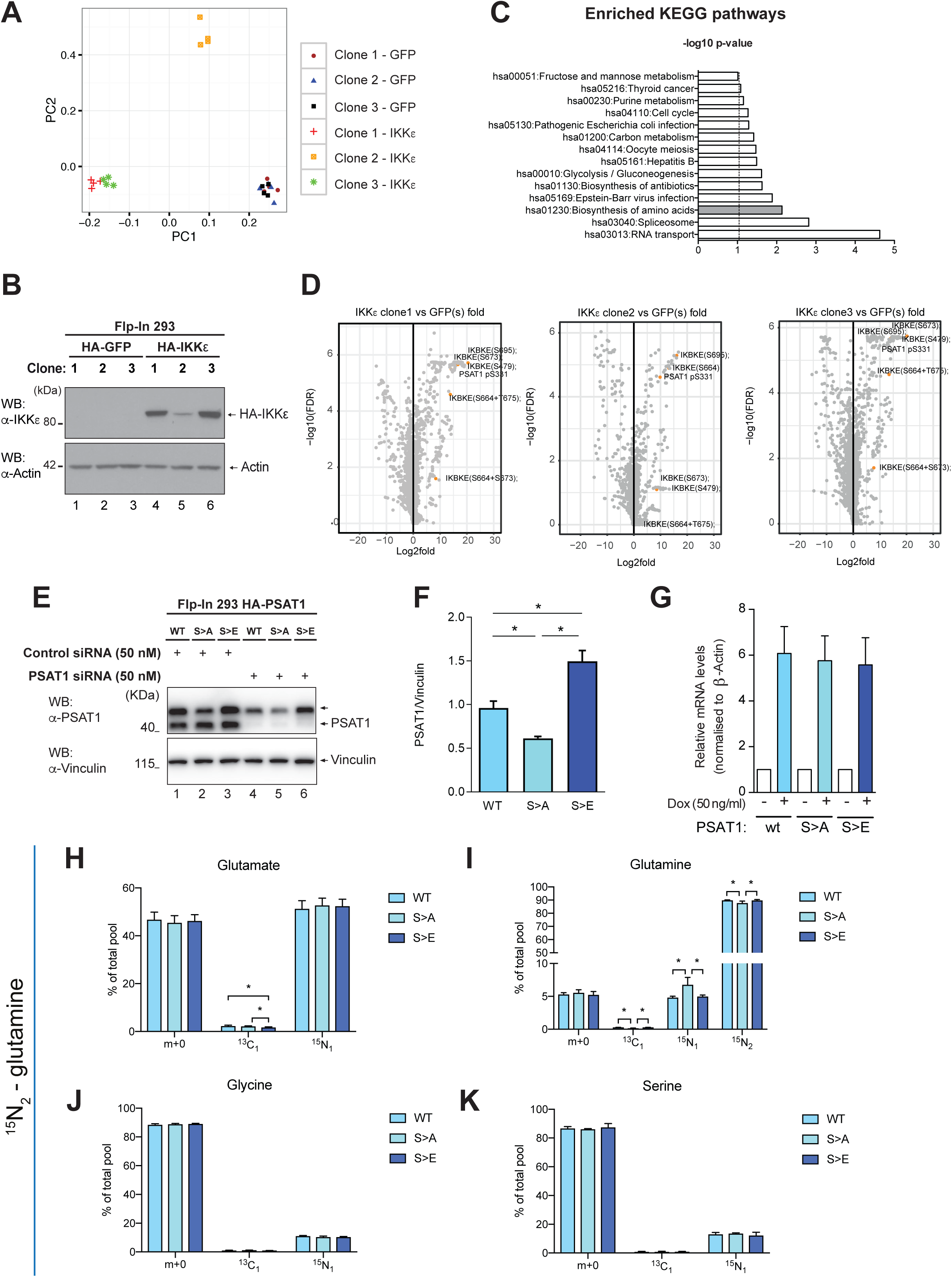
Direct phosphorylation of PSAT1 does not contribute to IKKε-induced serine biosynthesis. (A) Principal component analysis of differentially phosphorylated substrates in three independent single cell clones of Flp-In 293 HA-GFP or Flp-In 293 HA-IKKε cells treated with doxycycline (100 ng/ml, 16 hours). The phosphoproteomes in the three clones were analysed by mass spectrometry as described in material and methods. (B) Level of IKKε in three independent single cell clones of Flp-In 293 HA-GFP or Flp-In 293 HA-IKKε cells following treatment with doxycycline (100 ng/ml, 16 hours). (C) Pathway enrichment analysis of phosphoproteomic data identifying modification of proteins in key metabolic and signalling pathways as a result of IKKε expression. Enriched pathways from differentially phosphorylated substrates were determined using the David pathway enrichment analysis tool (see material and methods). (D) Volcano plot representation of statistically significant differentially phosphorylated substrates in indicated Flp-In 293 HA-IKKε expressing clones relative to HA-GFP expressing controls (n=4 independent replicates). P values were calculated by unpaired Student’s t-test of log transformed normalised phosphopeptide intensity values (peak areas of extracted ion chromatograms). Highlighted dots represent IKKε auto-phosphorylation sites indicative of kinase activity, and PSAT1 S331 phosphosite. (E) Level of endogenous PSAT1 and HA-tagged wild-type (WT) or mutant (S>A, S>E) PSAT1 in Flp-In 293 HA-PSAT1 wt, Flp-In 293 HA-PSAT1 S>A and Flp-In 293 HA-PSAT1 S>E cells treated with doxycycline (50 ng/ml, 16 hours). Endogenous PSAT1 was silenced in indicated samples prior to doxycycline treatment. (F) Levels of HA-tagged PSAT1 variants in endogenous *PSAT1*-silenced Flp-In 293 HA-PSAT1 wt, Flp-In 293 HA-PSAT1 S>A and Flp-In 293 HA-PSAT1 S>E cells treated with doxycycline (50 ng/ml, 16 hours). Densitometry analysis quantified single sample density as a percentage of total blot densitometry prior to vinculin normalisation (n=9, mean ± SEM, *p<0.05, one-way ANOVA with Bonferroni *post-hoc* tests). (G) Quantitative Real-Time PCR (qRT-PCR) analysis of *PSAT1* mRNA level in Flp-In 293 HA-PSAT1 wt, Flp-In 293 HA-PSAT1 S>A and Flp-In 293 HA-PSAT1 S>E cells treated with doxycycline (50 ng/ml, 16 hours). Data are expressed as relative to levels in non-treated cells and normalised to *β-Actin* (n=4, mean ± SEM) (H-K) ^15^N_2_-glutamine flux analysis of endogenous *PSAT1*-silenced Flp-In 293 HA-PSAT1 wt, Flp-In 293 HA-PSAT1 S>A and Flp-In 293 HA-PSAT1 S>E cells treated with doxycycline (50 ng/ml, 16 hours). Fractional enrichment of (H) glutamate, (I) glutamine, (J) glycine and (K) serine ^15^N-isotopologues. m+1 shows the naturally occurring ^13^C isotopologue (n=10). Data Information: In (E) and (H-K), endogenous PSAT1 was silenced by transfection with a pool of 3 siRNA oligos targeting the non-coding sequence of *PSAT1* mRNA. In (H-K), data are presented as mean ± SD, *p<0.05 (one-way ANOVA with Bonferroni *post-hoc* tests).

### Phosphorylation of PSAT1 on serine 331 does not affect its activity

To understand the functional consequences of PSAT1 phosphorylation, we generated a non-phosphorylatable (S331A, S>A) and a phospho-mimic (S331E, S>E) mutant enzyme via site-directed mutagenesis and expressed them in the inducible Flp-In 293 cell system. WB analysis showed significant, but rather small differences in the protein levels of the mutants, where PSAT1 S>A mutant was present at lower levels and the S>E mutant levels were elevated as compared to the WT protein (Supplementary Fig. S2A,B). Moreover, the alteration of the protein levels of the mutants was independent of the presence of endogenous PSAT1, excluding an effect on dimer formation by mutation of the enzyme (see Fig. 3E,F). No differences of *PSAT1* mRNA were observed between the cells expressing the mutants (Fig. 3G). Since differences in the level of expression of the mutants were modest, we tested whether the mutations had any effect on PSAT1 function. Analysis of steady state metabolite levels upon ^15^N_2_-glutamine labelling showed only minor differences in the percentage of labelled glycine and glutathione among the mutants (Fig. 3H-K). Thus, these experiments indicated that phosphorylation of PSAT1 does not alter its enzyme activity, and the observed small changes in abundance are not sufficient to induce serine biosynthesis.

As a further functional test, we also compared the proliferation rate of Flp-In 293 HA-PSAT1 cells expressing WT, S>A or S>E mutant PSAT1 while knocking-down endogenous PSAT1 via siRNA. As shown in Supplementary Fig. S2C-E, siRNA of PSAT1 or induction of PSAT1 wt, S>A or S>E did not affect the proliferation rate of cells growing in complete medium. Notably, this is different from the effect of PHGDH knock-down, which abolishes proliferation in a set of cell lines even in complete medium (Possemato *et al*, 2011). Importantly, however, we showed that PSAT1 knock-down impairs the ability of Flp-In 293 cells to grow in serine-free medium. The effect on proliferation was fully rescued by induction of PSAT1 wt or the S>E mutant, but the S>A mutant was not able to fully substitute endogenous PSAT1. Thus, while these experiments indicated that direct PSAT1 phosphorylation has functional consequences on cellular proliferation rate, it did not explain how IKKε induces serine biosynthesis.

### IKKε induces SBP gene transcription via a non-canonical target

In order to understand the primary mechanism of IKKε-induced serine biosynthesis, we assessed the overall effect of IKKε on PSAT1 protein levels, by measuring the abundance of the endogenous enzyme following induction of IKKε in Flp-In 293 cells and observed that IKKε induced a large accumulation of endogenous PSAT1 (Fig. 4A). Importantly, silencing of IKKε in a panel of breast cancer cell lines (ZR-75-1, T47D, MDA-MB-468, Cal120 and HCC1143) had the opposite effect, leading to a reduction in PSAT1 and in some cell lines also to decreased PHGDH and PSPH levels (Fig. 4B-E). Moreover, we observed that IKKε induction in Flp-In 293 HA-IKKε cells led to a marked (2-6 fold) increase in the transcription of all three SBP enzyme mRNAs (Fig. 4F). Again, this transcriptional effect was also confirmed in breast cancer cell lines, where upon siRNA-mediated silencing of IKKε *PSAT1* mRNA was reduced in 5 out of 9 cell lines (Fig. 4G). These results indicated that IKKε regulates the SBP primarily at the transcriptional level.

**Figure 4.**
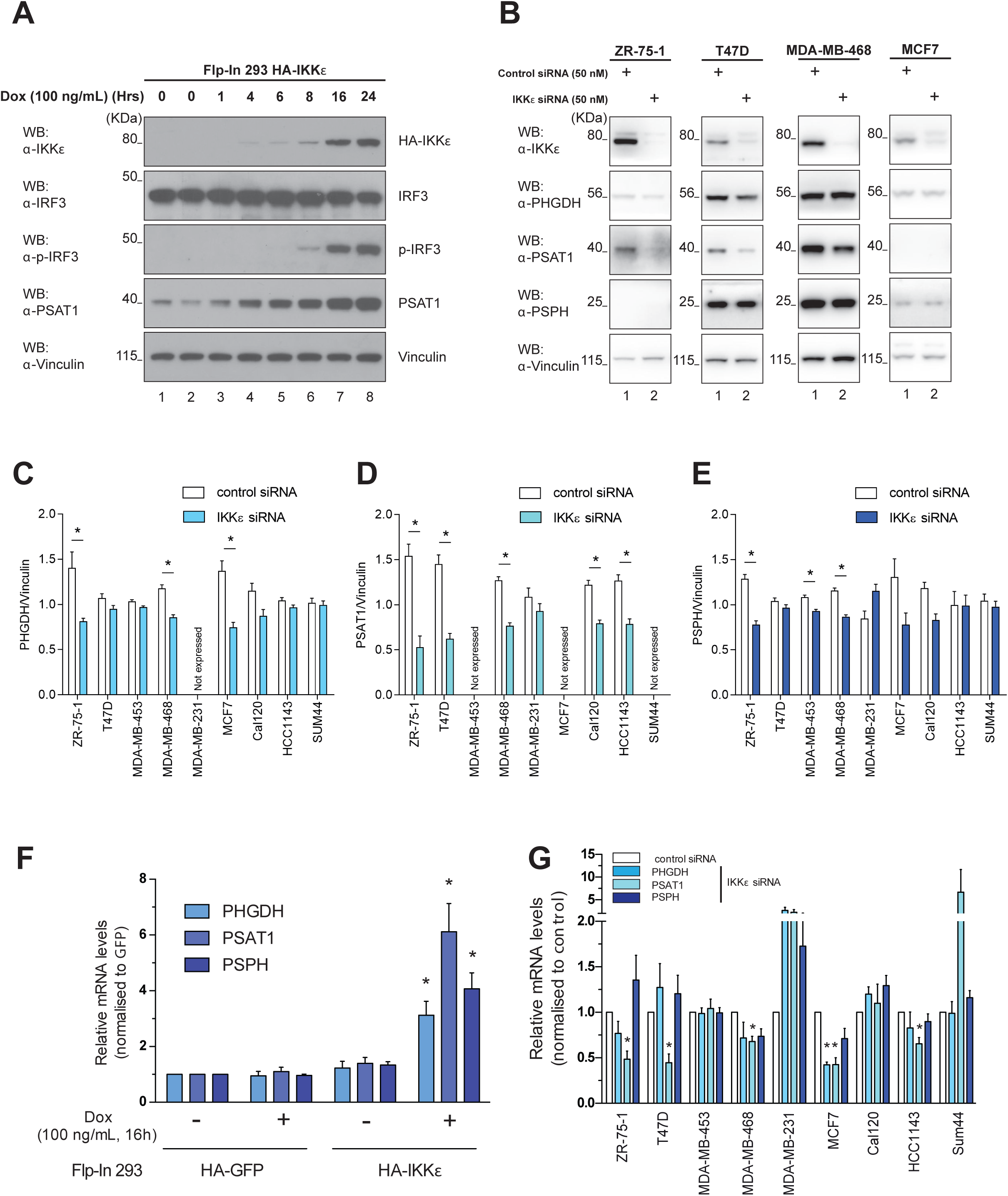
The SBP is primarily regulated by an IKKε-mediated transcriptional response. (A) Levels of IKKε, IRF3, phosphorylated IRF3 (S396) and PSAT1 in Flp-In 293 HA-IKKε cells treated with doxycycline (100 ng/ml) for the indicated number of hours. (B) Levels of IKKε, PHGDH, PSAT1 and PSPH in *IKBKE*-silenced ZR-75-1, T47D, MDA-MB-468 and MCF7 breast cancer cell lines. (C-E) Levels of the SBP enzymes in a panel of *IKBKE*-silenced breast cancer cell lines. (C) PHGDH, (D) PSAT1 and (E) PSPH level normalised to Vinculin. Densitometry analysis quantified single sample density as a percentage of total blot density per cell line prior to vinculin normalisation (n≥3, mean ± SEM, *p<0.05, two-tailed Student’s t-test). (F) qRT-PCR analysis of *PHGDH, PSAT1* and *PSPH* mRNA levels in Flp-In 293 HA-GFP or Flp-In 293 HA-IKKε cells treated with doxycycline (100 ng/ml, 16 hours). Data are expressed as relative to levels in non-treated Flp-In 293 HA-GFP cells and normalised to *β-Actin* (n=4, mean ± SEM, *p<0.05, two-way ANOVA with Bonferroni *post-hoc* tests) (G) qRT-PCR analysis of *PHGDH, PSAT1* and *PSPH* mRNA levels in a panel of *IKBKE*-silenced breast cancer cell lines. Data are expressed as relative to levels in a non-silenced control of each cell line and normalised to *β-Actin* (n≥3, mean ± SEM, *p < 0.05, as measured by one sample t-test).

To understand the mechanistic details of transcriptional induction of the SBP by IKKε, we first asked whether this effect requires the kinase activity of IKKε, and whether TBK1 is involved in the response. Silencing or knockout (via CRISPR-Cas9) of TBK1 had no effect on the IKKε-mediated induction of PSAT1 (Supplementary Fig. S3A-C). Moreover, we also observed that the kinase domain of IKKε was required to induce PSAT1 expression (Supplementary Fig. S3D). These results suggested that IKKε exerts its effect via up-regulating the enzymes of the pathway in a manner dependent from its kinase domain and independently from TBK1. Next, we tested the involvement of the canonical downstream signalling pathways known to be activated by the kinase. Silencing of the transcription factors IRF-3 and p65 (Clément *et al*, 2008) did not abolish the induction of SBP enzyme gene transcription by IKKε (Supplementary Fig. S3E-G). Of note, these experiments also indicated that IRF3 is required to maintain the transcription of the SBP genes in basal conditions, but has no role in IKKε-mediated induction. Finally, since IKKε also induces secretion of a range of cytokines (Barbie *et al*, 2014), we tested the possibility that the induction of SBP enzyme gene transcription is mediated by an autocrine loop. Extracellular media conditioned by GFP or IKKε expressing cells was collected and applied on three different receiving cell lines: Flp-In 293 HA-GFP, not expressing IKKε (Supplementary Fig. S4A); and the T47D and ZR-75-1 breast cancer cell lines, constitutively expressing IKKε (Supplementary Fig. S4B,C). Media conditioned by IKKε expressing cells had no effect on the SBP enzymes, even if cytokine-mediated signalling was observed in all three receiving cell lines, as demonstrated by induction of STAT1, phospho-STAT1 and OAS1 (an interferon inducible gene, only in ZR-75-1).

Altogether, these results demonstrated that IKKε kinase activity, independently of TBK1, induces SBP enzyme gene transcription by a cell-autonomous mechanism, which however does not include its canonical downstream targets.

### IKKε inhibits mitochondrial function via reducing PDH activity

Mitochondrial dysfunction has recently been proposed to cause up-regulation of the SBP (Meiser *et al*, 2016; Nikkanen *et al*, 2016). We therefore hypothesised that IKKε could alter mitochondrial function, as suggested by our initial experiments using tracer compounds (see Figs. 1 and 2), ultimately contributing to the transcriptional regulation of the SBP. Consistent with this hypothesis, mitochondrial oxygen consumption rate (OCR) was suppressed by IKKε induction in the Flp-In 293 HA-IKKε cell line (Fig. 5A), accompanied by reduced mitochondrial membrane potential (Δψ_m_, Fig. 5B), as assessed by respirometry and steady state tetramethyl-rhodamine methylester (TMRM) intensity imaging, respectively. Accordingly, IKKε silencing resulted in significantly higher OCR in a set of breast cancer cells (Fig. 5C) and this was not due to an off-target effect of the siRNAs used (Supplementary Fig S5A). In a set of control experiments we have shown that IKKε primarily affected ATP-coupled respiration, without significantly inhibiting uncoupled or reserve OCR, as shown by measuring respiration in the presence of oligomycin (inhibitor of the ATP synthase), and the uncoupler FCCP, respectively (Supplementary Fig. S5B-F). Moreover, the effect was integral to mitochondria, since mitochondrial isolated from IKKε expressing cells showed reduced respiration as compared to GFP-expressing controls (Supplementary Fig. S5G). Accordingly, this regulation was not mediated by PSAT1, since siRNA-mediated silencing of PSAT1 did not revert the IKKε-induced repression of mitochondrial function in either Flp-In 293 or in breast cancer cell lines (Fig. 5D, E).

**Figure 5.**
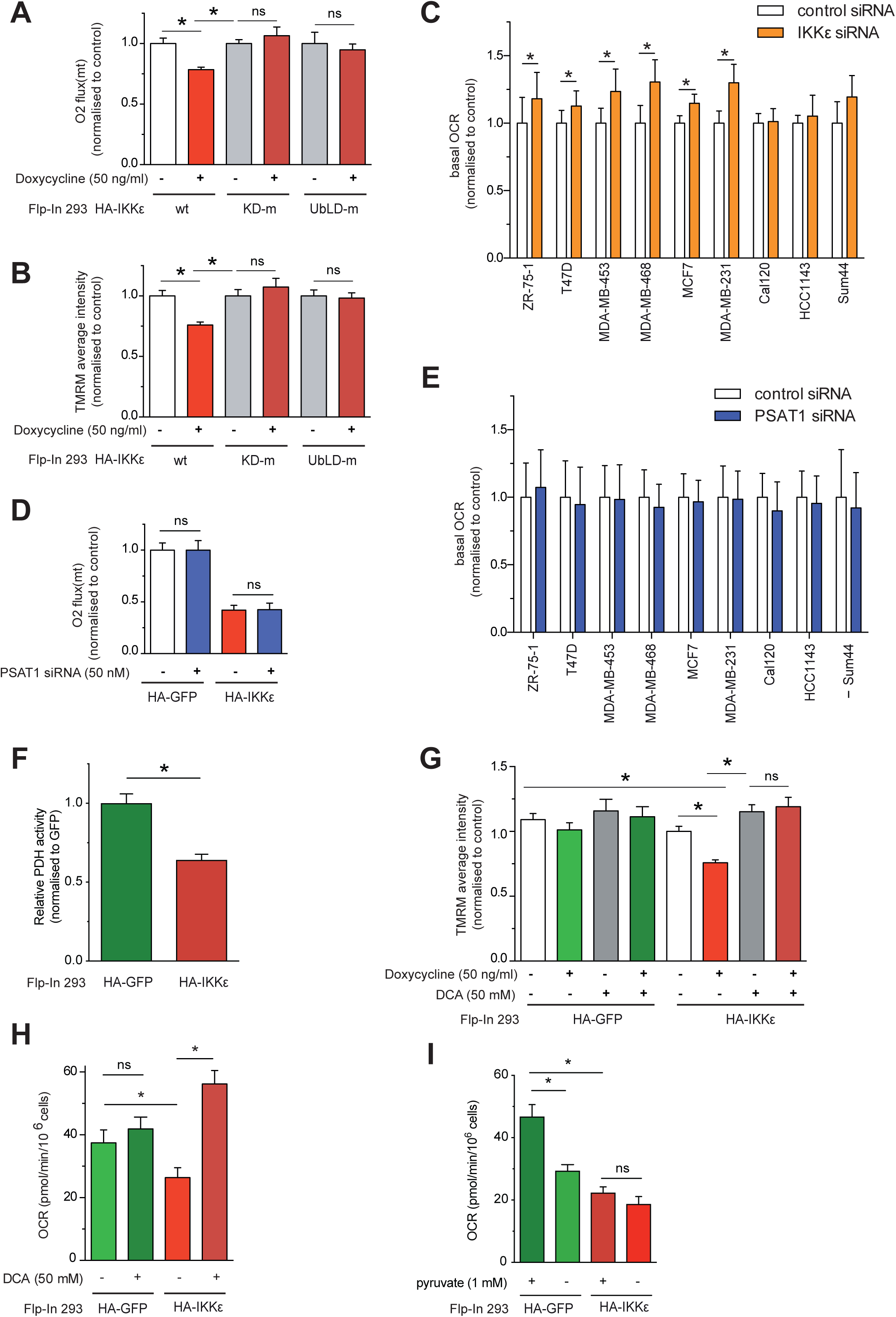
IKKε inhibits mitochondrial metabolism via reducing PDH activity. (A) Basal oxygen consumption in Flp-In 293 HA-IKKε wt, Flp-In 293 HA-IKKε KD-m and Flp-In 293 HA-IKKε UbLD-m cells following treatment with doxycycline (16 hours), measured using Oroboros high resolution respirometry. Data are normalised to non-treated control cells. (B) Average TMRM staining intensity in Flp-In 293 HA-IKKε wt, Flp-In 293 HA-IKKε KD-m and Flp-In 293 HA-IKKε UbLD-m expressing cells induced by doxycycline (16 hours). Data are normalised to non-treated controls cells. (C) Basal mitochondrial oxygen consumption rate (OCR) in a panel of *IKBKE*-silenced breast cancer cell lines, transfected with a pool of 4 targeting siRNA oligos, measured using Seahorse XF96 analysis (n ≥ 3, mean ± SEM, *p < 0.05, paired). (D) Basal oxygen consumption in *PSAT1-*silenced Flp-In 293 HA-GFP or Flp-In 293 HA-IKKε cells treated with doxycycline (16 hours). Cells were transfected with a pool of 4 targeting siRNA oligos. Measured using Oroboros high resolution respirometry. Data are normalised to non-treated Flp-In 293 HA-GFP control cells. (E) Basal OCR in a panel of *PSAT1-*silenced breast cancer cell lines, transfected with a pool of 4 targeting siRNA oligos, measured using Seahorse XF96 analysis. (n=3, mean ± SEM). (F) Relative pyruvate dehydrogenase (PDH) activity in Flp-In 293 HA-GFP or Flp-In 293 HA-IKKε cells treated with doxycycline (50 ng/ml, 16 hours) (n=3, mean ± SEM, *p < 0.05 by Student’s t-test). (G) Average TMRM staining intensity in Flp-In 293 HA-GFP or Flp-In 293 HA-IKKε cells treated with doxycycline and DCA (both for 16 hours). Data are normalised to non-treated Flp-In 293 HA-IKKε cells (n=3, mean ± SEM, *p < 0.05 as measured by two-way ANOVA with Bonferroni *post hoc* test). (H) Basal OCR in Flp-In 293 HA-GFP or Flp-In 293 HA-IKKε cells treated with doxycycline (50 ng/ml) in combination with dichloroacetate (DCA) for 16 hours, measured using Oroboros high resolution respirometry (n=8). (I) Basal OCR in Flp-In 293 HA-GFP or Flp-In 293 HA-IKKε cells treated with doxycycline (50 ng/ml) in combination with pyruvate deprivation for 16 hours, measured using Oroboros high resolution respirometry (n=8) Data Information: All data are n≥3 and presented as mean ± SEM, *p<0.05. In (A-D, H, I), data were normalised to total sample protein concentration. In (A, B) two-way ANOVA, in (G, H) one-way ANOVA with Fischer’s LSD *post-hoc* tests, in (C, E, F) two tailed paired Student’s t-tests for each cell line (C, E), and in (D, I) one-way ANOVA with Tukey’s multiple comparison tests were applied.

In our phosphoproteomic screen we showed that PDHA1 pS232 was significantly hyper-phosphorylated following IKKε induction (Table 2). Of note, other phosphosites were not changing or were less phosphorylated, indicating that the increase in pS232 is not due to a higher level of expression of the protein. This residue is known to inhibit PDH activity when phosphorylated and also reported to be necessary for tumour growth (Golias *et al*, 2016), thus we hypothesised that IKKε regulates pyruvate entry in the TCA cycle, and consequently electron provision for the respiratory chain. In agreement with this hypothesis, PDH activity was reduced in IKKε expressing cells (Fig. 5F). Thus, we tested the role of this phosphorylation on the metabolic changes triggered by IKKε by inhibiting pyruvate dehydrogenase kinases, responsible for PDH complex phosphorylation, using dichloroacetic acid (DCA) (Stacpoole, 1989). DCA reverted the IKKε-mediated reduction of Δψ_m_ and inhibition of respiration in Flp-In 293 mitochondria, but had no effect in control cells (Fig. 5G,H), indicating that diminished pyruvate oxidation by the PDH complex is the limiting factor of respiratory activity in IKKε overexpressing cells. In line with this conclusion, we also showed that IKKε overexpressing cells rely less on pyruvate for their respiration, in comparison to control cells, expressing GFP (Fig. 5I).

### IKKε activates an ATF4-mediated transcriptional response

Mitochondrial dysfunction was shown to regulate SBP gene transcription via Activating Transcription Factor 4 (ATF4) (Bao *et al*, 2016; Khan *et al*, 2017). Thus, we hypothesized that IKKε-induced mitochondrial inhibition elicits a similar ATF4-mediated response. Confirming this hypothesis, ATF4 was induced in IKKε expressing cells (Fig. 6A) and silencing of ATF4 abolished the transcriptional up-regulation of SBP enzymes by IKKε, and reduced their protein levels (Fig. 6B,C). Moreover, the same effect was seen in breast cancer cell lines upon the silencing of ATF4 (Fig. 6D,E), in agreement with previous data (DeNicola *et al*, 2015). Altogether, these data indicated that IKKε orchestrates a complex metabolic reprogramming that encompasses the inactivation of mitochondrial metabolism and the consequent transcriptional activation of the SBP, mediated by ATF4.

**Figure 6.**
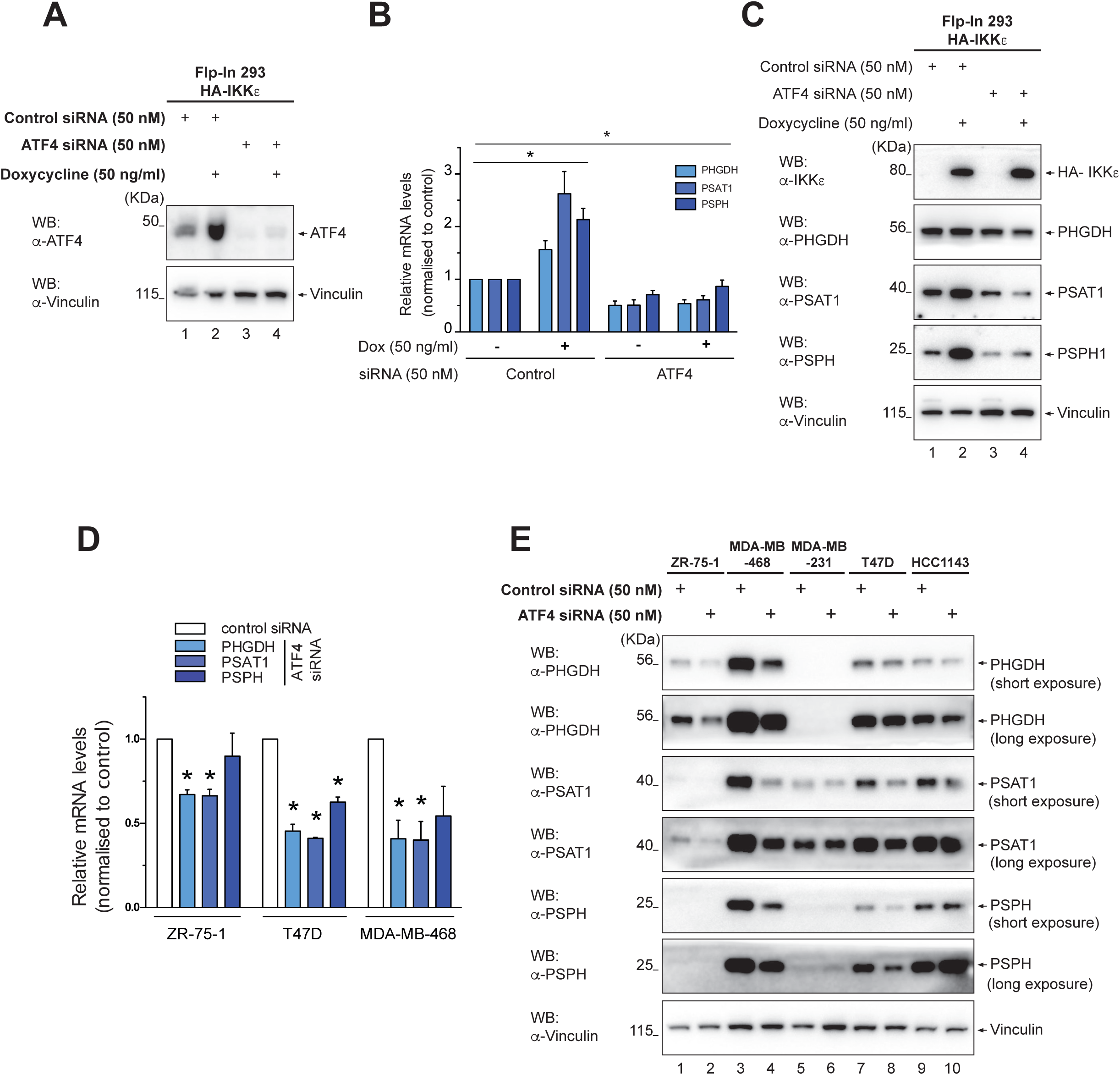
IKKε stimulates SBP enzyme gene transcription via ATF4. (A) Level of ATF4 in both non-silenced and *ATF4-*silenced Flp-In 293 HA-IKKε cells treated with doxycycline (16 hours). (B) qRT-PCR analysis of *PHGDH, PSAT1* and *PSPH* mRNA levels in both non-silenced and *ATF4-*silenced Flp-In 293 HA-IKKε cells treated with doxycycline (16 hours). Data are expressed as relative to levels in a non-silenced, non-treated control, and normalised to *β-Actin* (n=4, mean ± SEM, *p < 0.05, two-way ANOVA with Bonferroni *post-hoc* tests). (C) Levels of IKKε, PHGDH, PSAT1 and PSPH in both non-silenced and *ATF4*-silenced Flp-In 293 HA-IKKε cells treated with doxycycline (16 hours). (D) qRT-PCR analysis of *PHGDH, PSAT1* and *PSPH* mRNA levels in *ATF4-*silenced ZR-75-1, T47D and MDA-MB-468 breast cancer cell lines. Data are expressed as relative to levels in non-silenced control cells and normalised to *β-Actin* (n=3, mean ± SEM, *p < 0.05, one-sample t-test). (E) Levels of PHGDH, PSAT1 and PSPH in *ATF4*-silenced ZR-75-1, MDA-MB-468, MDA-MB-231, T47D and HCC1143 breast cancer cell lines.

### Pharmacological inhibition of IKKε-induced metabolic changes reduces cell proliferation

To test the functional consequences of the IKKε-mediated metabolic rewiring on tumour proliferation, we assessed the outcome of inhibiting the two key metabolic reactions of serine biosynthesis on cell proliferation in a panel of breast cancer cell lines. NCT502, a recently described inhibitor of the SBP (Pacold *et al*, 2016), but not its PHGDH-inactive form, significantly reduced breast cancer cell proliferation (Supplementary Fig. S6A,B). Importantly, the effect on proliferation was significantly correlated with the effect of IKKε on mitochondrial OCR, but not extracellular acidification rate (Fig. 7A and Supplementary S6C). These results suggested that inhibition of the specific metabolic effect regulated by IKKε correlates with cancer cell proliferation rate. The same effect was observed upon treatment with the glutamine antagonist 6-diazo-5-oxo-l-norleucine (DON) (Cervantes-Madrid *et al*, 2015) and CB839 (Gross *et al*, 2014), to inhibit glutaminase and thus glutamate availability for PSAT1 (Figs. 7B, Supplementary S6D-F). These results suggest that IKKε-induced metabolic changes promote cell proliferation and might be valuable targets to inhibit IKKε driven tumorigenesis.

**Figure 7.**
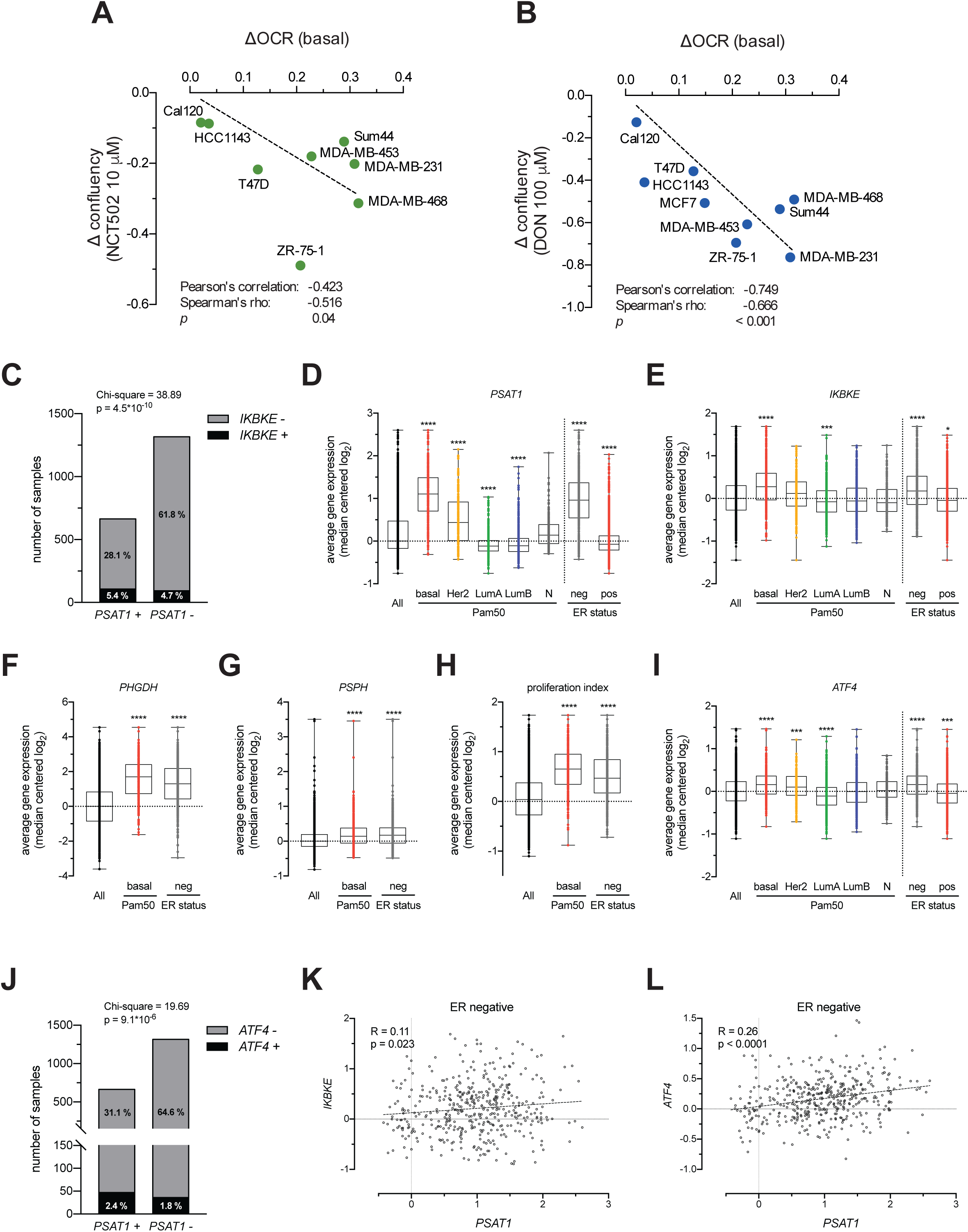
The pathophysiological role of IKKε-induced metabolic and gene expression alterations in breast cancer. (A) Correlation of change in OCR (ΔOCR) in a panel of *IKBKE*-silenced breast cancer cell lines and the change in cell confluency (Δconfluency) upon treatment of the panel of cell lines with NCT502. (B) Correlation of ΔOCR in a panel of *IKBKE*-silenced breast cancer cell lines and the Δconfluency upon treatment of the panel of cell lines with 6-Diazo-5-oxo-L-norleucine (DON). (C) Association between *IKBKE* and *PSAT1* mRNA overexpression evaluated by a chi-squared independence test. The + sign indicates samples with significant (p<0.05) overexpression of *IKBKE* or *PSAT1*. Number and percentage of samples, as well as the Chi-square values are shown. (D-I) Expression of *IKBKE* and the SBP enzymes *PSAT1*, *PHGDH* and *PSPH* in the METABRIC dataset. The expression of a proliferation related gene set and *ATF4* are also shown. Samples were stratified by Pam50 intrinsic subtypes and ER status. Ordinary one-way ANOVA and Tukey’s multiple comparison test were applied. *p<0.05, **p<0.02, ***p<0.002, ****p<0.001. (J) Association between *IKBKE* and *ATF4* mRNA overexpression evaluated by chi-squared independence test. The + sign indicates samples with significant (p<0.05) overexpression of *IKBKE* or *PSAT1*. Number and percentage of samples, as well as the Chi-square values are shown. (K, L) Correlation of *IKBKE*, *PSAT1* (K) and *ATF4* (L) expression in the ER negative sample subset. Data Information: In (A-B) *IKBKE-*silenced cells were transfected with a pool of 4 targeting oligos, cell confluency was measured using the IncuCyte Zoom. In (A, B, K, L) linear regression, correlation coefficients (Pearson’s correlation, Spearman’s rho) significance of difference from slope = 0 are shown. In (D-I) and (J) statistics are shown for all samples or for the first 50 samples of the rankings shown in (C-E).

### IKKε and PSAT1 is overexpressed in a common, highly proliferative subset of breast cancer

In order to explore IKKε and SBP enzyme gene expression status in breast cancer patient samples, we analysed data of the METABRIC dataset (Curtis *et al*, 2012), which includes data from 1981 breast cancer patients with pathological and clinical details. *IKBKE* and *PSAT1* mRNA were significantly up-regulated (above the 95% confidence interval) in 200 (10.1%) and 664 (33.5%) samples, respectively, and 107 (5.4%) samples showed overexpression of both mRNAs. This indicated a highly significant association between the two genes, as confirmed chi-square independence test (Fig. 7C). Given that breast cancer is a heterogenous disease, commonly classified into five to ten intrinsic subtypes (Perou *et al*, 2000; Curtis *et al*, 2012), the association of *IKBKE* and *PSAT1* might be driven by subtype specific expression. Thus, in order to identify subtypes with significant overexpression compared to the total population, we compared the expression values of *IKBKE* and the SBP genes in all Pam50 subtypes (Jiang *et al*, 2016) (Parker *et al*, 2009) and in the estrogen receptor (ER) positive and negative populations. This analysis indicated that both *IKBKE* and *PSAT1,* similarly to *PHGDH* and *PSPH*, are significantly upregulated in an ER negative Pam50:basal subpopulation of tumours, with the highest proliferation index (Nielsen *et al*, 2010) (Fig. 7D-H). ATF4 driven overexpression of *PSAT1* in ER negative tumours has been previously shown in a different dataset (Gao *et al*, 2017) and here we also have found strong association of these two genes (Fig. 7I, J). Importantly, while *PSAT1* is overexpressed in almost all ER negative samples (378 out of 435), only 79 samples overexpress *IKBKE*, indicating that *PSAT1* is regulated by multiple inputs. However, above 90% of samples overexpressing *IKBKE* (72 out of 79) also overexpress *PSAT1*. Due to the large fraction of PSAT1 overexpressing samples in the ER negative subset and the overall low expression and detected variability of *IKBKE* and *ATF4* mRNAs in the dataset, only low but still statistically significant levels of correlation could be found between these genes (Fig. 7K, L). However, these gene expression patterns clearly show that IKKε mediated expression of PSAT1 and the SBP enzymes, demonstrated in our in vitro experiments, is potentially also functional in a subset of clinical samples, suggesting the pathophysiological importance of the pathway in breast cancer.

## DISCUSSION

Here we described a novel fundamental mechanism by which IKKε, a key player in the innate immune response, directly regulates cellular metabolism. We show that the kinase orchestrates a complex metabolic reprogramming culminating in serine biosynthesis. The principal mechanism involved in IKKε-mediated regulation of the SBP is inhibition of carbon supply to the mitochondria, leading to the transcriptional up-regulation of SBP genes via a mitochondrial-nuclear retrograde signalling pathway targeting ATF4, ultimately activating serine biosynthesis. In parallel, IKKε, presumably in complex with its homolog TBK1, also phosphorylates PSAT1, regulating its expression level to a small degree. The overall metabolic switch induced by IKKε supports cancer cell proliferation and is present in a subset of breast tumours, providing potentially important pharmacological targets.

While such mechanistic details of the function of IKKε as newly identified modulator of the SBP and mitochondria have not been reported before, previous studies implicated IKKε in the regulation of cellular metabolism. Consistent with our data, IKKε was shown to inhibit OCR in MEFs (Reilly *et al*, 2013), and regulate glycolysis in DCs, although in this system the kinase did not affect mitochondrial metabolism (Everts *et al*, 2014). Similarly, in pancreatic ductal adenocarcinoma, IKKε was shown to stimulate glucose uptake, but did not inhibit mitochondrial respiration (Zubair *et al*, 2016). Thus, IKKε appears to modulate cellular metabolism in a tissue- and context-specific manner, and our study pinpoints and extends the breadth of the specific cellular targets utilized by the kinase to exert these heterogeneous responses. Importantly, in addition to the previously known canonical NFκB and IRF3 signalling pathways (Clément *et al*, 2008), IKKε can engage in metabolic regulation by direct or indirect phosphorylation of key metabolic enzymes (PSAT1 and PDH, respectively) and these modifications can exert a downstream effect on gene expression, by engaging mitochondrial-nuclear ATF4-mediated signalling. While here we have shown that in breast cancer cells IKKε-mediated changes in metabolism support proliferation, these metabolic alterations might also facilitate other cellular functions (Jones & Bianchi, 2015), for example, cytokine secretion in immune cells (Chang *et al*, 2013; Tannahill *et al*, 2013; Rodriguez *et al*, 2019; Yu *et al*, 2019). Apart from providing novel mechanistic details of IKKε-mediated cellular metabolic changes, this work also indicates the necessity of further research to better understand the physiological and pathological role of IKKε in order to efficiently and selectively target tumour cells relying on this oncogene. Our observations suggest that drugs targeting IKKε-regulated metabolic pathways can specifically target breast cancer cells without affecting other cell types, considering that only in these cells IKKε has been reported to regulate the SBP. Indeed, our gene expression analysis indicated that the IKKε-mediated pathway defines a subset of ER^-^, basal breast tumours, thus evaluation of the IKKε-mediated metabolic and gene expression phenotype can help to further stratify breast cancer for treatment. While more investigation is required to elucidate the functional consequence of IKKε-induced PSAT1 phosphorylation, analysis of the recent phospho-proteomic dataset of breast cancer (Mertins *et al*, 2016) also indicated that PSAT1 phosphorylation is associated with triple negative, basal type (see Table 3), and our stratification is also in agreement with previous studies, where strong correlation between *PSAT1* expression and tumour proliferation has been found in ER^-^ tumours (Coloff *et al*, 2016; Gao *et al*, 2017). Of note, IKKε, along with the JAK/STAT pathway, has been reported to regulate a cytokine network promoting cellular proliferation in a subset of triple negative breast tumours (Barbie *et al*, 2014), and PSAT1 overexpression is also a feature of a small fraction of ER^+^ tumours along with the JAK-STAT pathway (De Marchi *et al*, 2017). While we have shown that IKKε activates the JAK-STAT pathway (see Supplementary Fig. S4) along with PSAT1 overexpression, JAK-STAT activation *per se* was not sufficient for the induction of the enzyme. This indicates that both IKKε and PSAT1 are defining features in order to identify tumours where the pathway is actively promoting proliferation.

Finally, further work is required to investigate if the IKKε-mediated metabolic phenotype described here, especially regarding the regulation of the SBP, occurs in different cellular systems where IKKε is known to be activated. This could help to develop new therapeutic strategies applicable in a broad range of inflammation-related diseases, beyond cancer.

## MATERIALS AND METHODS

### Plasmids

DNA fragments encoding human IKKɛ (UniProt: Q14164) or PSAT1 (UniProt: Q9Y617) were amplified separately by PCR using primers containing Kpn1 (5’) and EcoR1 (3’) restriction sites. The PCR products were double digested by these two enzymes and ligated to vector pCDNA5.5 which provides a 2xHA tag at the c-terminal (a kind gift from Dr Tencho Tenev). *UbLD-M-IKKɛ* plasmid was a kind gift of Prof Ivan Dikic. The other mutant variants of IKKε and PSAT1 were created by site directed mutagenesis and then ligated to pCDNA5.5 using the same method as the wide type.

### Cells

Flp-In 293 cells (Invitrogen, UK) were transfected with either *pcDNA5.5-wt-IKKɛ*, *pcDNA5.5-KD-M-IKKɛ* (K38A), *pcDNA5.5-UbLD-M-IKKɛ* (L353A F354A), *pcDNA 5.5-wt-PSAT1*, *pcDNA5.5-S331A-PSAT1*, *pCDNA5.5-S331E-PSAT1* or *pcDNA5.5-GFP,* together with a pOG44 plasmid at a molar ratio of 1:9. Stable cell lines and single cell clones were selected with 300 µg/ml hygromycin (Calbiochem, UK). All Flp-In 293 cells were cultured in DMEM (Sigma, UK).The panel of breast cancer cell lines was kindly provided by Dr. Alice Shia and Prof. Peter Schmid. MDA-MB-231, MDA-MB-468, MDA-MB-175, ZR75.1, T47D, HCC1143, MCF7 were cultured in RPMI-1640 (Sigma), Cal120 and MDA-MB-453 were cultured in DMEM (Sigma) and Sum44 in DMEM (Sigma) and 1 nM estrogen (Sigma). For all cell lines, medium was supplemented with 10% FBS, Penicillin-Streptomycin and Normocin (InvivoGeN). Serine free medium was custom made DMEM without serine, with 10% dialyzed FBS and Penicillin-Streptomycin. All cells were cultured with environmental conditions of 37°C, 5% CO_2_.

### siRNA

AllStars Negative Control siRNA (1027281, Qiagen, UK); Hs_ATF4_9 FlexiTube siRNA (SI04236337, Qiagen, UK); Hs_IKBKE_6 FlexiTube siRNA, (S102622319, Qiagen, UK); Hs_IKBKE_7 FlexiTube siRNA (S102622326); Hs_IKBKE_8 FlexiTube siRNA (S102655317, Qiagen, UK); Hs_IKBKE_9 FlexiTube siRNA (s102655324, Qiagen, UK); Hs_IRF3_4 FlexiTube siRNA (SI02626526, Qiagen, UK); Hs_PSAT1_10 FlexiTube siRNA (SI03019709, Qiagen, UK); Hs_PSAT1_12 FlexiTube siRNA (SI03222142, Qiagen, UK); Hs_PSAT1_14 FlexiTube siRNA (SI04265625, Qiagen, UK); Hs_PSAT1_15 FlexiTube siRNA (SI04272212, Qiagen, UK); Hs_RELA_5 FlexiTube siRNA (SI00301672, Qiagen, UK); Hs_TBK1_1 FlexiTube siRNA (SI00100961, Qiagen, UK); Hs_TBK1_5 FlexiTube siRNA (SI00301889, Qiagen, UK); Hs_TBK1_6 FlexiTube siRNA (SI02224411, Qiagen UK); Hs_TBK1_7 FlexiTube siRNA (SI02224418, Qiagen, UK) All the siRNA were used at a final concentration of 50 nM.

### Primers

**For IKKɛ ligation:**

For 5’-ttggtaccagccagctcagggcaggagatgcagagcacagccaatta-3’

Rev 5’-gatggatatctgcagaattcaggaggtgctgggactc-3’

### For PSAT1 ligation

For 5’-ttggtaccagccatggacgcccccaggcag-3’

Rev 5’-gatggatatctgcagaattctagctgatgcatctcc-3’

**For IKKɛ K38A mutagenesis:**

For 5’-gagctggttgctgtggcggtcttcaacactac-3’

Rev 5’-gtagtgttgaagaccgccacagcaaccagc-3’

**For PSAT1 S331A mutagenesis:**

For 5’-gaactcaatatgttggccttgaaagggc-3’

Rev 5’-gccctttcaaggccaacatattgag-3’

**For PSAT1 S331E mutagenesis:**

For 5’-gaactcaatatgttggacttgaaagggc-3’

Rev 5’-gccctttcaagtccaacatattgag-3’

### TaqMan™ Gene Expression Probes

*PHGDH* (Hs00198333_m1, ThermoFisher), *PSAT1* (Hs00795278_mH, ThermoFisher), *PSPH* (Hs00190154_m1, ThermoFisher), *ACTB* (β-Actin)(4310881E, Applied Biosystems).

### Drugs

6-Diazo-5-oxo-L-norleucine (Don, D2141, Sigma, UK); Sodium dichloroacetate (DCA, 347795, Sigma, UK); NCT-502 and PHGDH inactive (19716 and 19717, Cayman, UK); Doxycycline (Dox, D9891, Sigma, UK), Oligomycin (sc-203342, Santa Cruz Biotechnology, UK); FCCP (sc-203578, Santa Cruz Biotechnology, UK); Antimycin (sc-202467, Santa Cruz Biotechnology, UK); Rotenone (sc-203242, Santa Cruz Biotechnology, UK); CB-839 (Focus Biomolecules); MG-132 (Sigma, UK); Lactacystin (Enzo Life Sciences) and Chloroquine (Sigma, UK); Adenosine diphosphate (ADP, A2754, Sigma, UK).

### Experimental Procedures

#### Oxygen consumption and extracellular acidification rate measurements

An XF24 Extracellular or XF96e Extracellular Flux analyzer (Seahorse Biosciences, Agilent Technologies, US) was used to determine the bioenergetic profiles in breast cancer cell lines. Cells were plated in 6-well corning dishes first, then transfected with siRNA 24h after plating. 24h after transfection, cells were trypsinized, counted and plated into a 24 or 96-well Seahorse plate. Oxygen consumption rates (OCR) and extracellular acidification rates (ECAR) were assessed in Seahorse medium according to the manufacturer protocols. Respiratory parameters were assessed as described in Fig S6A. Oxygen consumption rate (OCR) of Flp-In 293 cells was measured using an Oroboros high resolution respirometer (Oroboros, Austria) at 37°C, in Seahorse XF assay medium containing 4.5 g/l glucose, 1 mM pyruvate and 25mM Hepes and the assay was performed as in Fig. S6A.

For measurements in isolated mitochondria, Flp-In 293 cells were first washed with PBS and collected in homogenization buffer (250 mM sucrose, 5 mM Hepes, pH 7.4, 0.5 mM EGTA, and Protease inhibitor cocktail (1187358001, Roche), then homogenized in a glass/glass, tight potter by 100 strokes on ice, followed by centrifugation for 5 min at 800xg at 4°C. The supernatant, containing mitochondria, was centrifuged again at 9000xg. The pellet was resuspended and adjusted to a protein concentration of 0.8 mg/ml in OCB buffer (125 mM KCl, 20 mM MOPS, 10 mM Tris ph7.2-7.3, 0.2 mM EGTA, 2.5 mM KH_2_PO_4_, 2.5 mM MgCl_2_). 10 mM glutamate and 5 mM malate were added to the mitochondrial suspension before the experiment, and OCR was measured in OCB buffer using the Oroboros high resolution respirometer. ADP (final concentration 0.25 mM), Cyt.C (10 µM - C2037, Sigma), oligomycin (5 µM) were injected step by step, and 50 µM FCCP was added in 1 µl steps until maximum respiratory capacity was detected. At the end of the run antimycin (5 µM final concentration) was injected. Data were the analyzed by the Datalab 5.5 (Oroboros, Austria) software.

### Cell proliferation assay

Cells were plated in Corning 96 well plates at a density between 2000 to 10,000 cells per well for different cell lines. Cell proliferation was then measured using the IncuCyte ZOOM instrument (Essen Biosciences, Ann Arbor, MI, USA) for 3-7d, and proliferation was analyzed with the Incucyte Zoom 2015A software.

### Metabolic labeling and metabolome analysis

Flp-In 293 cells and breast cancer cell lines (T47D and MDA-MB-468) were first plated separately in 6-well plates in five technical replicas per each condition. IKKε expression in Flp-In 293 cells was then induced by 50 ng/ml doxycycline and breast cancer cells were transfected with siRNA to suppress IKKɛ. 2h after induction for the Flp-In 293 cells, and 48h after siRNA transfection for the breast cancer cell lines, cells were incubated with either ^13^C_6_-glucose (CLM-1396-5, Cambridge Isotope Laboratories) or ^15^N_2_-glutamine (NLM-1328-0.25, Cambridge Isotope Laboratories) for 14h. Cells were then washed three times with PBS, and metabolites were extracted using cold extraction buffer (50% methanol, 30% acetonitrile, 20% ultrapure water, 100 ng/mL HEPES) at a ratio of 1ml extraction buffer/10^6^ cells. After 15 min incubation time on methanol and dry ice, cells were placed on a shaker for 15 min using a thermal mixer at 4°C, and incubated for 1h at −20°C. Cell lysates were centrifuged and the supernatant was collected and transferred into autosampler glass vials, which were stored at −80°C until further analysis.

Samples were randomized in order to avoid bias due to machine drift and processed blindly. LC-MS analysis was performed using a Q Exactive Hybrid Quadrupole-Orbitrap mass spectrometer coupled to a Dionex U3000 UHPLC system (Thermo Fisher Scientific). The liquid chromatography system was fitted with a Sequant ZIC-pHILIC column (150 mm × 2.1 mm) and guard column (20 mm × 2.1 mm) from Merck Millipore (Germany) and temperature maintained at 45°C. The mobile phase was composed of 20 mM ammonium carbonate and 0.1% ammonium hydroxide in water (solvent A), and acetonitrile (solvent B). The flow rate was set at 200 µL/min with the gradient described previously (Mackay *et al*, 2015). The mass spectrometer was operated in full MS and polarity switching mode. The acquired spectra were analyzed using XCalibur Qual Browser and XCalibur Quan Browser software (Thermo Scientific).

### Phosphoproteomics

#### Sample preparation

Flp-In 293 single cell clones for IKKɛ (Clone 1,2 and 3) and GFP (Clone 1,2 and 3) were seeded in 6-well plate in three replica for each condition. After 24h of seeding, cells were induced with doxycycline for 16h. Cells were first washed with ice cold PBS containing 1 mM Na_3_VO_4_ and 1 mM NaF, and then lysed in a lysis buffer containing 8M Urea, 20 mM HEPES, 1 mM Na_3_VO_4_, 1mM NaF, 1mM B-Glycerol phosphatate and 0.25 mM Na_2_H_2_P_2_O_7_. After incubation on ice for 5min, the cells were then scraped and collected in Eppendorf tubes and stored at −80°C. For sample analysis, cell lysates were thawed, protein digested with trypsin, and phosphopeptides were enriched using TiO_2_ as described in (Wilkes and Cutillas, 2017).

#### Nanoflow-liquid chromatography tandem mass spectrometry (LC–MS/MS)

Dried samples were dissolved in 0.1% TFA (0.5µg/µl) and run in a LTQ-Orbitrap XL mass spectrometer (Thermo Fisher Scientific) connected to a nanoflow ultra-high pressure liquid chromatography (UPLC, NanoAcquity, Waters). Peptides were separated using a 75 µm × 150 mm column (BEH130 C18, 1.7 µm Waters) using solvent A (0.1% FA in LC– MS grade water) and solvent B (0.1% FA in LC–MS grade ACN) as mobile phases. The UPLC settings consisted of a sample loading flow rate of 2 µL/min for 8 min followed by a gradient elution with starting with 5% of solvent B and ramping up to 35% over 100 min followed by a 10 min wash at 85% B and a 15 min equilibration step at 1% B. The flow rate for the sample run was 300 nL/min with an operating back pressure of about 3800 psi. Full scan survey spectra (m/z 375–1800) were acquired in the Orbitrap with a resolution of 30000 at m/z 400. A data dependent analysis (DDA) was employed in which the five most abundant multiply charged ions present in the survey spectrum were automatically mass-selected, fragmented by collision-induced dissociation (normalized collision energy 35%) and analysed in the LTQ. Dynamic exclusion was enabled with the exclusion list restricted to 500 entries, exclusion duration of 30 s and mass window of 10 ppm.

#### Database search for peptide/protein identification and MS data analysis

Peptide identification was by searchers against the SwissProt database (version 2013-2014) restricted to human entries using the Mascot search engine (v 2.5.0, Matrix Science, London, UK). The parameters included trypsin as digestion enzyme with up to two missed cleavages permitted, carbamidomethyl (C) as a fixed modification and Pyro-glu (*N*-term), Oxidation (M) and Phospho (STY) as variable modifications. Datasets were searched with a mass tolerance of ±5 ppm and a fragment mass tolerance of ±0.8 Da.

The automated program Pescal (Cutillas & Vanhaesebroeck, 2007) was used to calculate the peak areas of the peptides identified by the mascot search engine. Proteins were identified with at least two peptides matched to the protein and a mascot score cut-off of 50 was used to filter false-positive detection. The resulting quantitative data were parsed into Excel files for normalization and statistical analysis. Significance was assessed by T-Test of log2 transformed data. When required, p-values were adjusted using the Benjamini-Hochberg method. Results are shown as log2 fold IKKε over control.

### Western Blot

For western blot, cells were lysed in a lysis buffer (20 mM Tris.HCl, pH 7.4, 135 mM NaCl, 1.5 mM MgCl2, 1% Triton, 10 % glycerol) containing cOmplete protease inhibitor cocktail (Roche) and HALT phosphatase inhibitor cocktail (78428, Thermo scientific). Samples were quantified using DC protein assay kit (BioRad) and equal concentration samples were then prepared for SDS-PAGE in loading buffer (40% Glycerol, 30% β-Mercaptoethanol, 6% SDS, bromophenol blue). SDS-PAGE was performed using either 10% or 4-12% NuPAGE™ Bis-Tris Protein gels (Invitrogen) and resolved protein was transferred to Immobilon-P PVDF 0.45µm Membrane (Merck). Proteins were detected using primary antibodies to Actin (Santa Cruz, sc-1615), HA-tag (Roche, 11867423001) IKKɛ (Sigma, 14907) IRF3 (Cell Signaling, 11904) p-IRF3 Ser396 (Cell Signaling, 4947), OAS1 (sc-98424, Santa Cruz), PHGDH (Sigma, HPA021241), PSAT1 (Proteintech Europe, 20180-1-AP), PSPH (Proteintech Europe, 14513-1-AP), P65 (Cell Signaling, 8242), p-P65 (Cell Signaling, 3039), STAT1 (Cell Signaling, 9172), p-STAT1 Tyr701 (Cell Signaling, 9167), TBK1 (Cell Signaling, 3013) and Vinculin (Proteintech Europe, 66305-1-Ig) and ATF4 (ab1371, Abcam).

### High content imaging and measurement of mitochondrial membrane potential (Δψ_m_)

Cells were seeded in thin, clear bottom black 96 well plates (BD Falcon) at medium density (4000 cells/well) 24 hours before the experiments. Prior to imaging cells were loaded with 1µg/mL Hoechst 33342 (Sigma, UK) and 30 nM tetramethyl-rhodamine-methylester (TMRM) for 30 min. TMRM was present during imaging in the solution (DMEM w/o phenol red). Images were acquired with the ImageXpress Micro XL (Molecular Devices) high content wide field digital imaging system using a Lumencor SOLA light engine illumination, ex377/50 nm em447/60 nm (Hoechst) or ex562/40 nm and ex624/40 nm (TMRM) filters, and a 60X, S PlanFluor ELWD 0.70 NA air objective, using laser based autofocusing. 16 fields/well were acquired. Images were analysed with the granularity analysis module in the MetaXpress 6.2 software (Molecular Devices) to find mitochondrial (TMRM) and nuclear (Hoechst) objects with local thresholding. Average TMRM intensities per cell were measured and averaged for each well. The mean of wells were then used as individual data for statistical analysis to compare each condition.

### PDH activity measurement

PDH activity was measured on whole cell lysates using the Pyruvate dehydrogenase (PDH) Enzyme Activity Microplate Assay Kit (ab109902, Abcam).

### Transfection

siRNA were mixed with Darmafect 4 (Dharmacon), and cells were transfected according to the manufacturer’s instruction for 48 h or 72 h prior to measurements. Unless otherwise stated, a pool of all 4 IKBKE-targeting oligos was used for suppression of IKKε, a pool of all 4 TBK1-targeting oligos was used for suppression of TBK1, a pool of all 4 PSAT1-targeting oligos was used for general suppression of PSAT1, a pool of 3 PSAT1-targeting oligos (12, 14 and 15) was used for specific suppression of endogenous PSAT1, and single targeting oligos were used for the suppression of ATF4, p65 and IRF3. cDNA plasmids were mixed with Lipofectamine LTX (15338100, Thermo Fisher Scientific) or Fugene HD (E2311, Promega) according to the manufacturer’s instruction and transfected into the different cell lines for 48 h.

### qPCR

Total RNA was extracted from cells using the RNeasy Mini Kit (Qiagen) as per the manufacturer’s protocol. RNA yield was quantified using the NanoDrop ND-1000 (Thermo Fisher) and 1 mg of RNA was reverse transcribed to cDNA using the Omniscript RT Kit (Qiagen). qPCR was performed using the TaqMan assay system. Assay mixtures were prepared consisting of 10 µl TaqMan Master Mix (Applied Biosystems), 1 µl TaqMan gene probe & 1 µl cDNA, topped up to 20 µl with 8 µl RNase free H_2_O. The qPCR reaction was carried out using either the 7500 Real Time or the QuantStudio 5 Real Time PCR systems (Applied Biosystems) and the process was 2 minutes at 50°C, followed by 10 minutes holding at 95°C, then 40 cycles of 15 seconds at 95°C and 1 minute at 60°C. Relative mRNA quantifications were obtained using the comparative Ct method, and data was analysed using either the 7500 software v2.3 or QuantStudio Design & Analysis Software (Applied Biosystems).

### *In vitro* kinase assay

Flp-In 293 cells expressing HA-IKKε were washed with ice cold PBS containing Na_3_VO_4_ and NaF phosphatase inhibitors and frozen at −80°C. Following thawing, cells were lysed on ice for 30 minutes in a kinase assay lysis buffer (20mM Tris HCl (pH 7.8), 135mM NaCl, 1.5mM MgCl_2_, 1% Triton, 10% Glycerol, 1 complete protease inhibitor cocktail tablet, 1mM Na_3_VO_4_, 2.5mM Na_2_P_2_H_2_O_7_, 20mM β-glycerol phosphate and 15nM Okadaic acid). HA-tagged IKKε was isolated and purified from 1mg of whole cell lysate by pull-down using Anti-HA Agarose beads (Sigma Aldrich) for 1hr at 4°C with continuous mixing. 5 pmol of purified recombinant PSAT1 protein (PSAT1-1328H, Creative Biomart) was added to the pelleted and washed agarose beads along with kinase buffer (25mM Tris HCl (pH 7.8), 10mM MgCl_2_, 0.5 mM EGTA, 1mM DTT, 1mM Na_3_VO_4_, 2.5mM Na_2_P_2_H_2_O_7_ and 20mM β-glycerol phosphate) and 10mM ATP (New England Biolabs) to a total volume of 50µl. The reaction was incubated at room temperature for 20 minutes and halted by the addition of solid urea to a final concentration of 8M.

Phosphorylation of recombinant PSAT1 was analyzed after proteolytic digestion into peptides using trypsin and peptides were desalted using C18+carbon top tips (Glygen corparation, TT2MC18.96) and eluted with 70% acetonitrile (ACN) with 0.1% formaic acid. 2 µl peptides were loaded to nanoflow EASY-Spray (ES803) ultimate 3000 RSL nano instrument coupled on-line to a Q Exactive plus mass spectrometer (Thermo Fisher Scientific). Gradient elution was from 3% to 35% buffer B in 60min at a flow rate 250 nL/min with buffer A being used to balance the mobile phase (buffer A was 0.1% formic acid in water and B was 0.1% formic acid in acetonitrile). The mass spectrometer was controlled by Xcalibur software (version 4.0) and operated in the positive mode. The spray voltage was 2 kV and the capillary temperature was set to 255°C. The Q-Exactive plus was operated in data dependent mode with one survey MS scan followed by 15 MS/MS scans. The full scans were acquired in the mass analyser at 375-1500m/z with the resolution of 70 000, and the MS/MS scans were obtained with a resolution of 17 500.

For quantification of phosphorylated and un-phosphorylated PSAT derived peptides, the extracted ion chromatograms were integrated using the theorectical masses of ions with 4 significant figures with mass tolerance of 5 ppm. Values of area-under-the-curvewere obtainded manually in Qual browser of Xcalibur software (version 4.0)

### Immunoprecipitation

Flp-In 293 cells expressing HA-IKKε or HA-PSAT1 were washed with ice cold and frozen at −80°C. Following thawing, cells were lysed on ice for 30 minutes in Triton 1% lysis buffer (20 mM Tris.HCl, pH 7.4, 135 mM NaCl, 1.5 mM MgCl_2_, 1% Triton, 10 % glycerol and 1 complete protease inhibitor cocktail tablet (Roche)). HA-tagged IKKε or PSAT1 were isolated and purified from 1mg of whole cell lysate by pull-down using Anti-HA Agarose beads (Sigma Aldrich) for 1hr at 4°C with continuous mixing. Beads were centrifuged at 4500rpm for 30 seconds and washed with lysis buffer 3 times before being resuspended in 0.2M glycine (pH 2.5) for 30 minutes to elute bound protein from the beads. Beads were centrifuged once more and glycine eluates were transferred to fresh tubes and pH was equilibrated with 1M ammonium bicarbonate. Loading dye was added to 1x concentration and samples were used for SDS-PAGE and western blotting.

### Generation of Conditioned Medium

Flp-In 293 HA-GFP or HA-IKKε cells were treated for 16 hours with 50 ng/ml doxycycline in 1 ml of medium per well of a 6 well plate, allowing secretion of potential signaling factors into the medium. Following induction, medium was collected and filtered using a 0.22 µM pore size filter, and stored at 4°C till use.

### Gene expression analysis of clinical samples

The METABRIC dataset (Curtis et al, 2012) was obtained from Synapse: http://www.synapse.org (syn1688369/METABRIC Data for Use in Independent Research). All analysis was carried out using Bioconductor R packages. Overexpression of all genes were determined by fitting a Gaussian distribution to the central subpopulation shifted to zero, and then determining samples which had expressions greater than 1.96 times the standard deviation from zero.

### CRISPR cell line generation

2.5 million Flp-In 293 HA-IKKε wt or KM cells were plated in a 10 cm diameter culture dish (Corning). The following day, cells were transfected with 4.5 μg of either pSpCas9(BB)-2A-GFP control plasmid DNA (Ran *et al*, 2013) or one of 2 different TBK1 targeting plasmids using Lipofectamine® LTX with Plus™ Reagent (Invitrogen). Briefly, 2 transfection mixes were prepared per culture dish. Mix A, consisting of 900 μl OptiMEM, 4.5 μg plasmid DNA and 22.5 μl Plus^TM^ Reagent, and mix B, consisting of 900 μl OptiMEM and 27 μl Lipofectamine® LTX were prepared independently, incubated for 5 minutes at room temperature, then mixed by pipetting and brief vortexing. The combined AB solution was then incubated for 30 minutes at room temperature before being added dropwise to the existing cell culture media in each dish. Following overnight incubation at room temperature, the culture media was changed to fresh media and the transfected cells were incubated for 48 hours, after which cells were sorted for positive GFP expression using BD FACSAriaII. GFP positive single cells were plated in individual wells of a 96 well plate. 12 cells were plated per plasmid/cell line combination and allowed to grow to single cell colonies. Growing colonies were then expanded into 6 well plates and checked for TBK1 knockout by western blotting.

## ACKNOWLEDGMENTS

We are grateful to Dr. Alice Shia and Prof. Peter Schmid (BCI – QMUL) for providing the breast cancer cell line panel and to Dr. Tencho Tenev and Prof. Pascal Meier (ICR, London) for useful advice and discussion, to Dr Gunnel Hallden (BCI – QMUL) for providing reagents, to Prof. Ivan Dickic (Institute of Biochemistry II – Goethe University, Germany) for providing plasmids containing IKKε mutants and to the Barts Cancer Institute FACS facility for their support. KB is supported by the Barts London Charity (Grant Reference Number: 467/2053) and WJ is supported by the Medical Research Council (MRC) PhD program. PC is funded by the BBSRC (BB/M006174/1) and Barts and The London Charity (297/2249). GS is funded by University College London COMPLeX/British Heart Foundation Fund (SP/08/004), the BBSRC (BB/L020874/1) and the Wellcome Trust (097815/Z/11/Z) in the UK, and the Italian Association for Cancer Research (AIRC, IG22221). CF and ASHC are funded by the Medical Research Council, core fund to the MRC Cancer Unit.

## AUTHORS CONTRIBUTION

RX performed experiments to characterise the effect of IKKε on mitochondrial oxygen consumption rate, and to test the sensitivity of breast cancer cell lines to different drugs. WJ performed the experiments to characterise the mechanism through which IKKε regulates cellular metabolism (RT-PCR and WB). RX and WJ did the cloning of IKKε mutants and PSAT1 mutants (respectively) and generated the cell lines. EWV helped with the phospho-proteomic experiment together with VR and PC, that also performed the MS for the in vitro kinase assay. ASHC and CF performed the MS experiment with metabolic tracers and analysed the data. AN and CC helped with phospho-proteomic data analysis. BY contributed to the OCR measurements, SOB to the characterization of the role of ATF4 and YW to the cloning. GS performed the experiments to measure mitochondrial membrane potential and analysed the data. RBB, GS and Kevin Bryson analysed the gene expression datasets. Katiuscia Bianchi designed the study and wrote the manuscript with the help of RX, WJ, PC, GS and CF.

---

**Table 1.** ^13^C_6_-glucose and ^15^N_2_-glutamine labelling data in Flp-In 293 HA-GFP vs HA-IKKε, T47D and MDA-MB-468 breast cancer cell lines upon transfection with control siRNA and IKKε siRNA.

**Table 2.** Phospho-proteomic data analysis

**Table 3.** Phosphorylation of S331 – PSAT1 in breast cancer (Mertins *et al*, 2016)

**Table 4.** Characteristics of all METABRIC samples in the mitochondria bicluster. The table contains the ranking, PC1 value, *IKBKE* and *PSAT1* expression values, average expression values in mitochondrial group1 and 2, as well as the GO terms identified for genes differentially expressed between the UF and LF samples.

## SUPPLEMENTARY FIGURE LEGENDS

**Figure S1.**
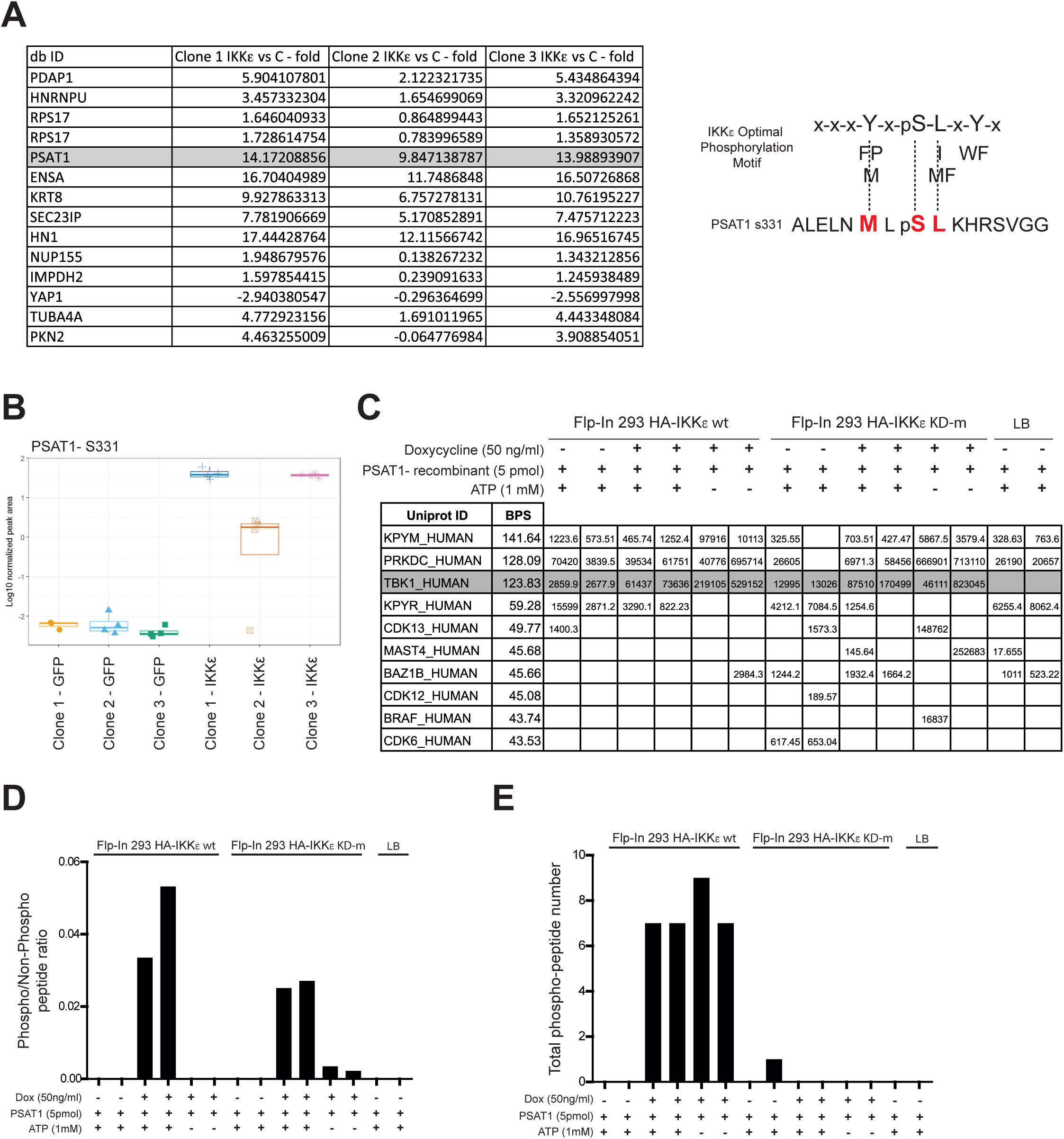
*In vitro* phosphorylation of PSAT1 by IKKε. (A) Left table: List of potential direct substrates for IKKε as estimated by motif of the identified phosphopeptides with high log2 fold changes. The high scoring PSAT1 is highlighted. Right panel: Overlap of PSAT1 S331 phosphosite sequence with 3 out of the 4 defined residues of IKKε’s recognition motif. (B) PSAT1 S331 phosphorylation in three independent single cell clones of Flp-In 293 HA-GFP and Flp-In 293 HA-IKKε cells treated with doxycycline (100 ng/ml, 16 hours). Quantified by MS-MS phosphopeptide intensity ratios. (C) Relative abundance of secondary kinases in *in vitro* kinase assay samples as detected by mass spectrometry analysis. (D) Phosphorylation of recombinant PSAT1 at S331 in an *in vitro* kinase reaction containing ATP and HA-IKKε wt, HA-IKKε KD-m or a blank lysis buffer control (LB). HA-tagged kinase variants were purified from corresponding Flp-In 293 cells treated with doxycycline (50 ng/ml, 16 hours). PSAT1 S331 phosphorylation was measured by mass spectrometry analysis. Data are normalised as a ratio of PSAT1 phosphopeptide to total levels. (E) The number of IKKε phosphopeptides detected in each *in vitro* kinase assay sample by mass spectrometry analysis.

**Figure S2.**
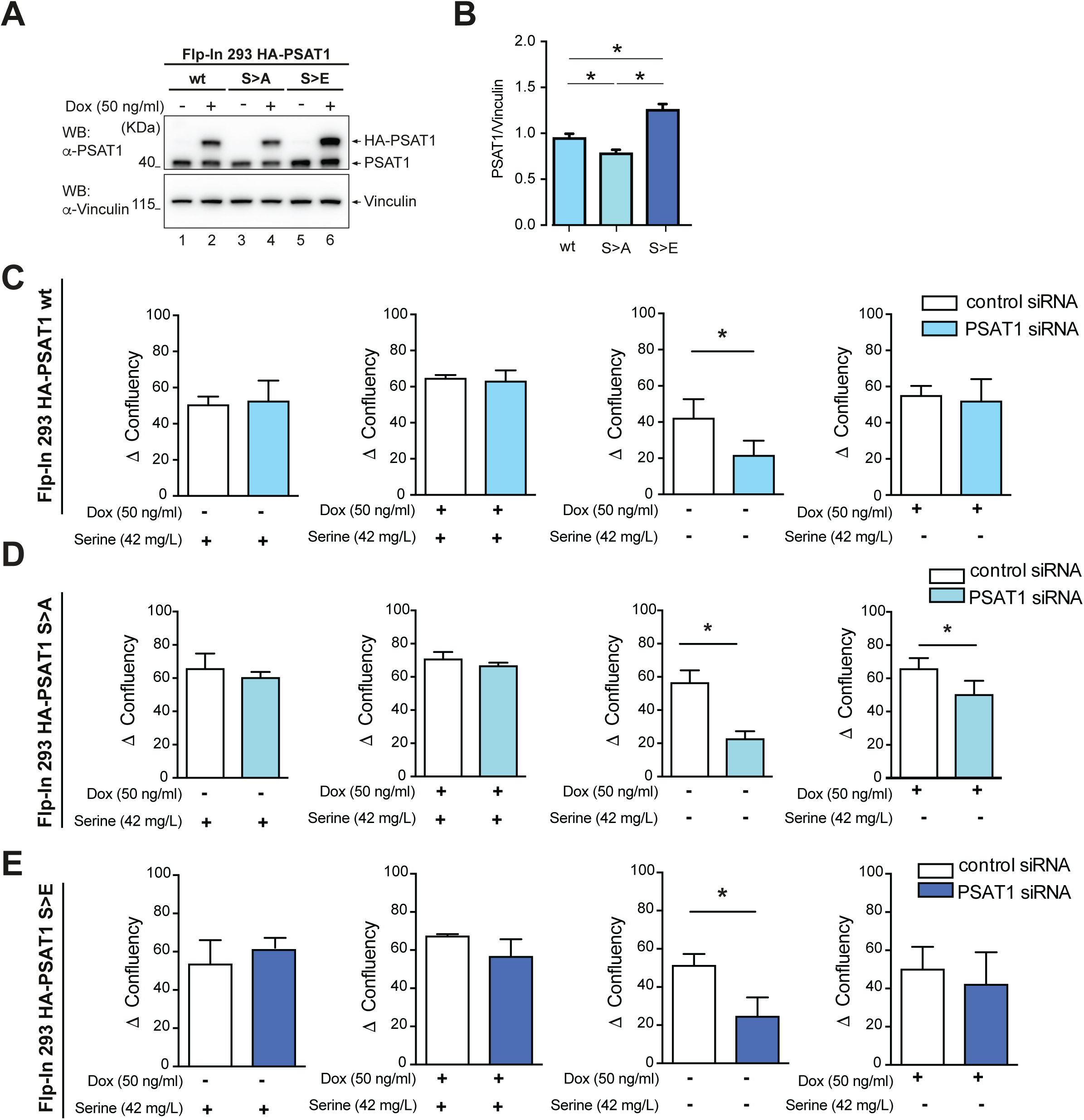
The effect of the PSAT1 S331 phosphomutants on cellular proliferation in response to serine deprivation. (A) Level of endogenous and HA-tagged PSAT1 in Flp-In 293 HA-PSAT1 wt, Flp-In 293 HA- PSAT1 S>A and Flp-In 293 HA-PSAT1 S>E cells treated with doxycycline (16 hours). (B) Levels of HA-tagged PSAT1 variants in endogenous *PSAT1-*silenced Flp-In 293 HA- PSAT1 wt, Flp-In 293 HA-PSAT1 S>A and Flp-In 293 HA-PSAT1 S>E cells treated with doxycycline (50 ng/ml, 16 hours). Densitometry analysis quantified single sample density as a percentage of total blot densitometry prior to vinculin normalisation (n=9, mean ± SEM, *p<0.05, one-way ANOVA with Bonferroni post-hoc tests). (C-E) IncuCyte Zoom analysis of change in confluency (Δ confluency) of (C) Flp-In 293 HA- PSAT1 wt, (D) Flp-In 293 HA-PSAT1 S>A and (E) Flp-In 293 HA-PSAT1 S>E cells over 72 hours. Both non-silenced and *PSAT1*-silenced cells were treated with doxycycline and grown in serine-free media or serine-free media reconstituted with serine as indicated. Data are expressed as mean ± SEM, n=3, *p<0.05 (paired students t-test). Data Information: In (B-E), endogenous *PSAT1-*silenced cells were transfected with a pool of 3 siRNA oligos targeting the non-coding regions of *PSAT1* mRNA.

**Figure S3.**
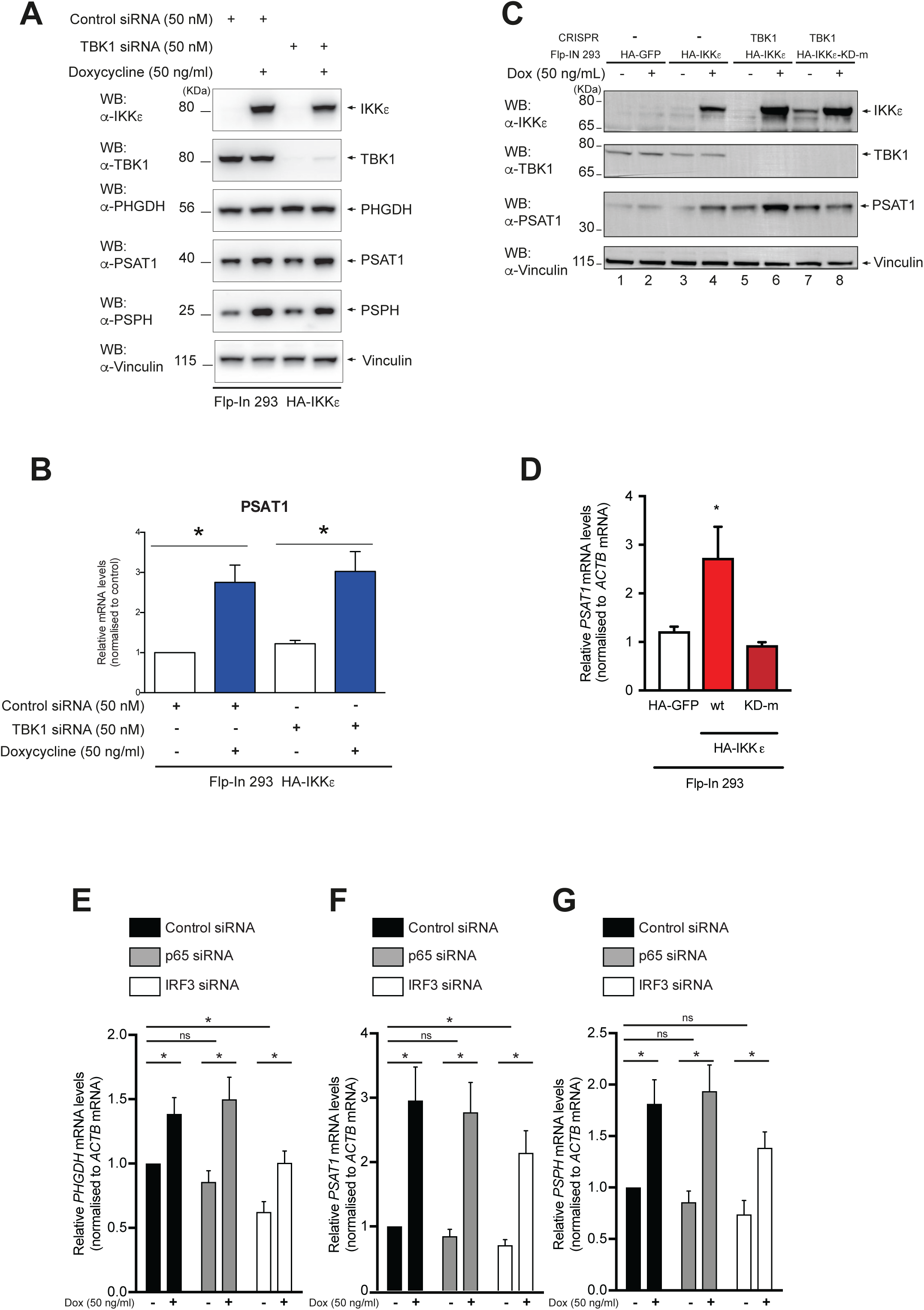
The effect of IKKε on SBP gene transcription is independent of TBK1, requires the IKKε kinase domain, and mediated by a non-canonical pathway. (A) Levels of IKKε, TBK1, PHGDH, PSAT1 and PSPH in both non-silenced and *TBK1*- silenced Flp-In 293 HA-IKKε cells, treated with doxycycline (16 hours). (B) qRT-PCR analysis of *PSAT1* mRNA levels in both non-silenced and *TBK1*-silenced Flp-In 293 HA-IKKε cells, treated with doxycycline (16 hours). Data are expressed as relative to levels in non-silenced, non-treated control cells and normalised to *β-Actin* (n=3, mean ± SEM, *p<0.05, unpaired t-test). (C) Levels of IKKε, TBK1 and PSAT1 in Flp-In 293 HA-GFP and Flp-In 293 HA-IKKε cells and TBK1 knockout Flp-In 293 HA-IKKε wt and Flp-In 293 HA-IKKε KD-m cells, generated using CRISPR Cas9 technology, following treatment with doxycycline (16 hours). (D) qRT-PCR analysis of *PSAT1* mRNA levels in Flp-In 293 HA-GFP, Flp-In 293 HA-IKKε wt and Flp-In 293 HA-IKKε KD-m cells treated with doxycycline (50 ng/ml, 16 hours). Data are expressed as relative to non-treated control cells and normalised to *β-Actin* (n=3, mean ± SEM, *p<0.05, two-way ANOVA with Bonferroni *post-hoc* tests). (E-G) qRT-PCR analysis of (E) *PHGDH*, (F) *PSAT1* and (G) *PSPH* mRNA levels in *p65-* or *IRF3*-silenced Flp-In 293 HA-IKKε cells treated with doxycycline (16 hours). Data are expressed relative to levels in non-silenced, non-treated cells and normalised to *β-Actin* (n≥4, mean ± SEM, *p<0.05, unpaired t-test). Data Information: In (A-B), *TBK1*-silenced cells were transfected with a pool of 4 targeting siRNA oligos. In (E-G), *p65-* and *IRF3-*silenced cells were transfected with single targeting oligos.

**Figure S4.**
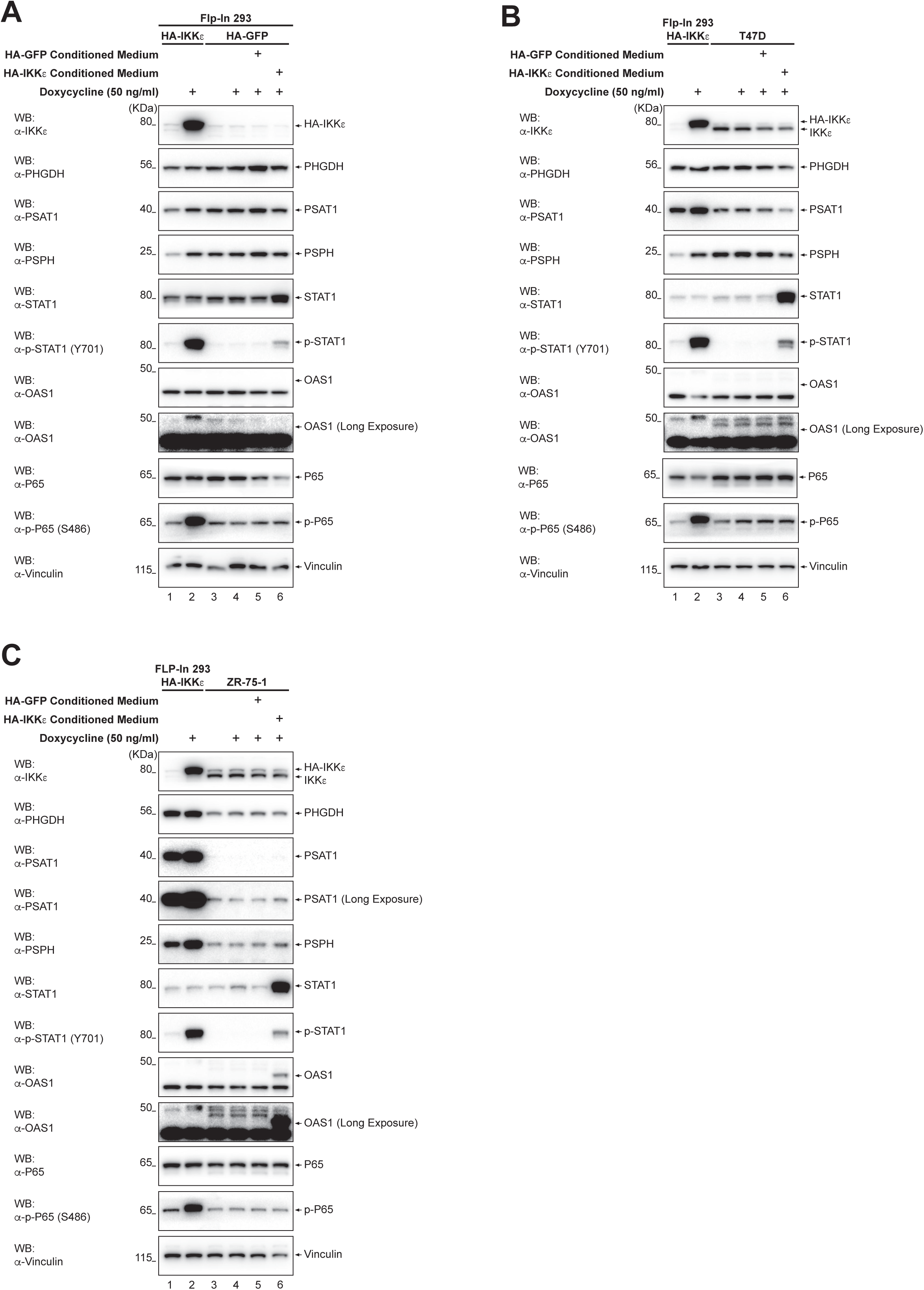
IKKε mediates cytokine secretion but no autocrine effect is responsible for the induction of SBP enzymes. (A-C) Levels of IKKε, PHGDH, PSAT1, PSPH, STAT1, phosphorylated STAT1 (Y701), OAS1, p65 and phosphorylated p65 (S486) in (A) Flp-In 293 HA-GFP cells, (B) T47D breast cancer cells and (C) ZR-75-1 breast cancer cells treated for 24 hours with media conditioned by Flp-In 293 HA-GFP or Flp-In 293 HA-IKKε cells treated with doxycycline (50 ng/ml, 16 hours). Changes in protein expression were compared to changes in Flp-In 293 HA-IKKε cells treated with doxycycline (16 hours).

**Figure S5.**
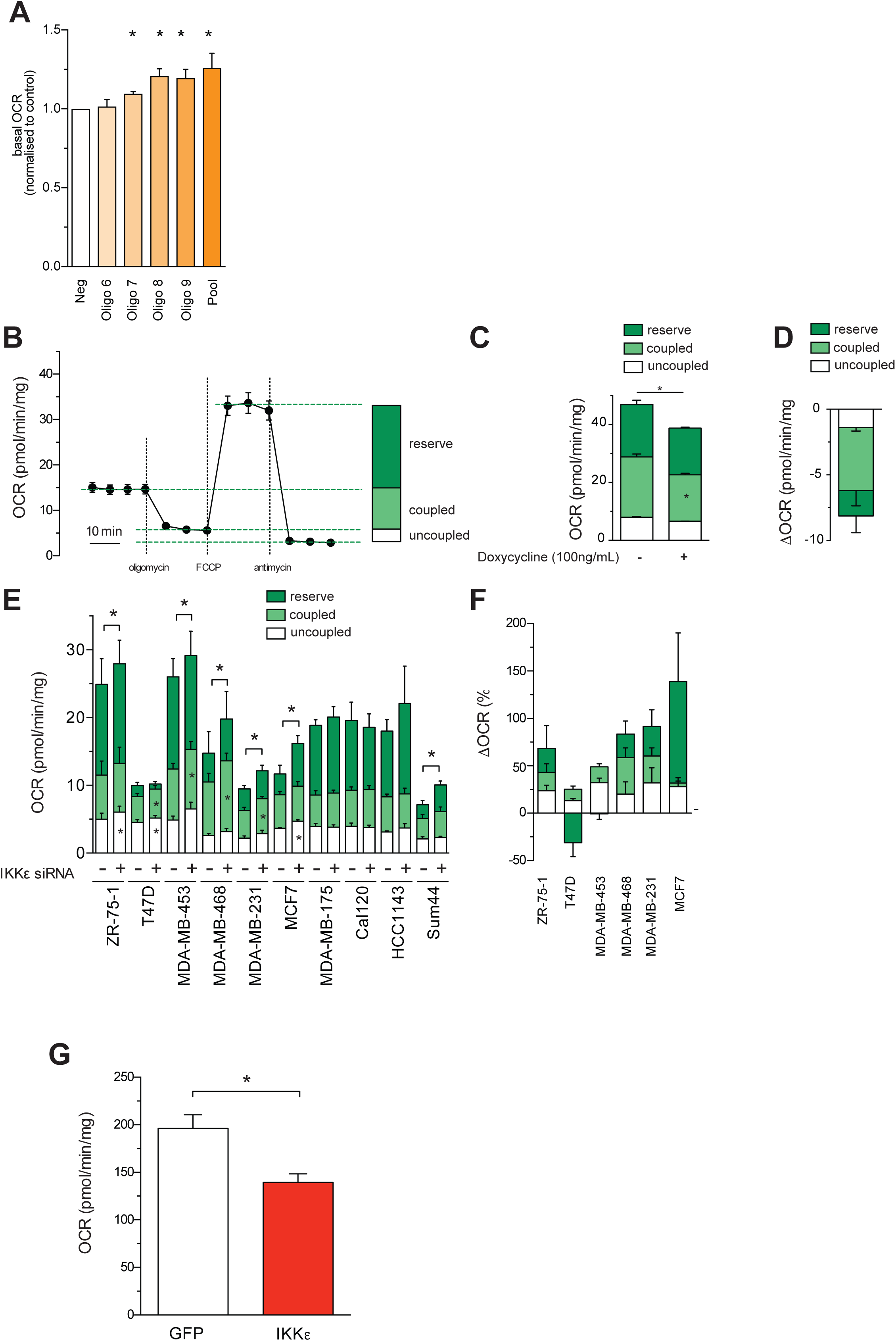
The effect of IKKε on mitochondrial respiration and membrane potential. (A) Effect of individual *IKBKE* siRNA oligos. Basal OCR in *IKBKE-*silenced MDA-MB-468 cells. Data are normalised to non-silenced control cells (Neg). (B) Schematic breakdown of Seahorse XF96 analysis, generating OCR curves showing different mitochondrial respiratory functions. Seahorse XF96 analysis determines basal, coupled, uncoupled and reserve respiratory capacities, allowing detailed characterisation of mitochondrial respiratory profiles in various treatment conditions (See Materials and Methods). (C) Reserve, coupled and uncoupled respiratory capacity profile of Flp-In 293 HA-IKKε cells. (D) Statistics of relative individual differences in reserve, coupled and uncoupled respiration in Flp-In 293 HA-IKKε cells. (E) Reserve, coupled and uncoupled respiratory capacity profiles of a selection of *IKBKE*- silenced breast cancer cell lines, where IKKε had a statistically significant effect on basal OCR (see Fig. 5C). (F) Statistics of relative individual differences in reserve, coupled and uncoupled respiration in indicated *IKBKE-*silenced breast cancer cells. (G) OCR in mitochondria isolated from Flp-In 293 HA-GFP or HA-IKKε cells treated with doxycycline (50 ng/ml, 16 hours). Data Information: Data were collected using Seahorse XF96 analysis (A, E, F) or with Oroboros (C, D, G), n ≥ 3, *p < 0.05. In (A, E, F) cells were transfected with a pool of 4 targeting siRNA oligos, or 1 of 4 targeting siRNA oligos that comprise the pool (A). In (C-D, G) cells were treated with doxycycline (100 ng/ml, 16 hours). In (A) one-way ANOVA with Fisher’s LSD test, in (C, E) two-way ANOVA with Bonferroni *post-hoc* tests, in (G) paired, two-tailed Student’s t-tests were applied.

**Figure S6.**
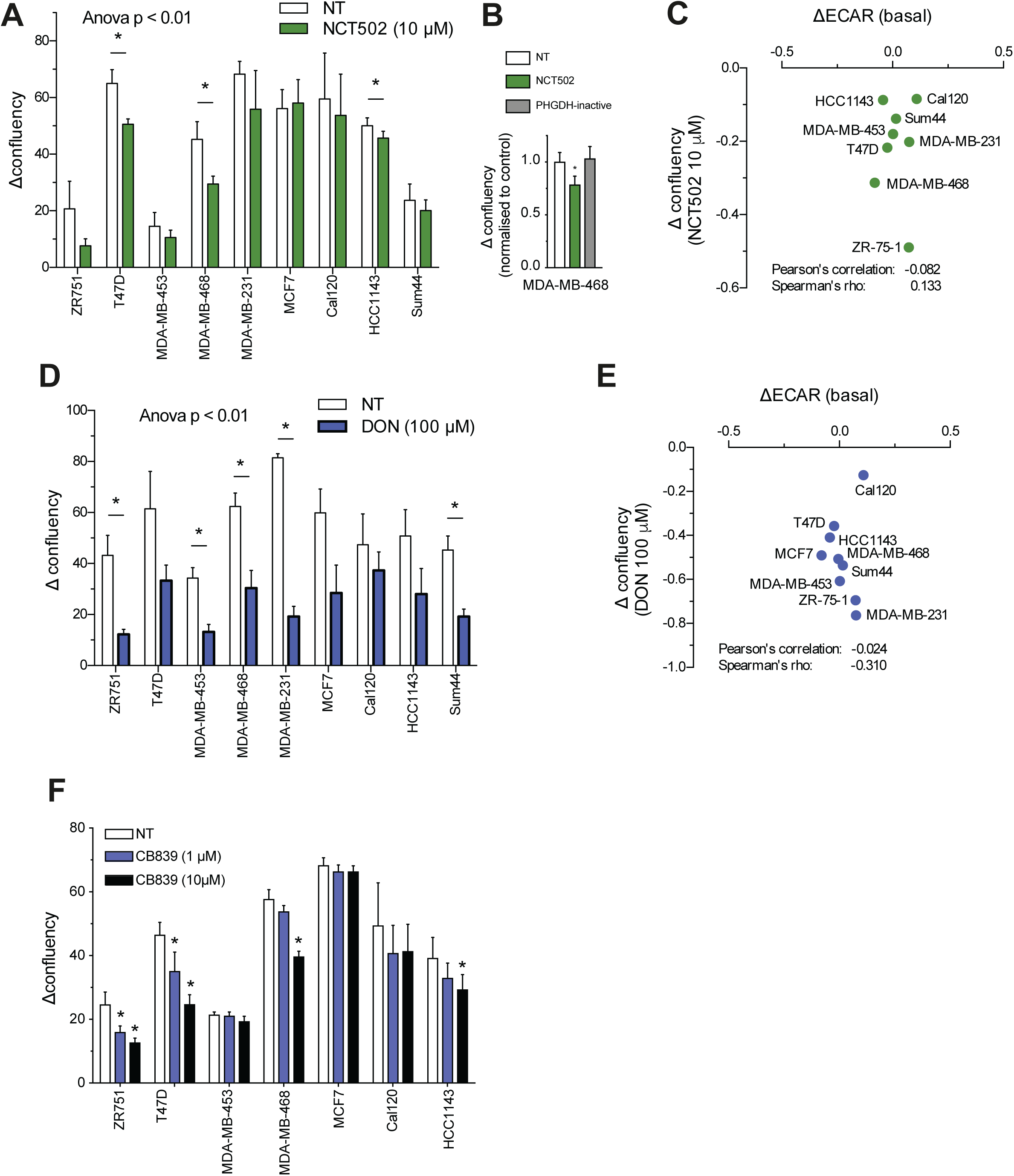
The pathophysiological role of IKKε-associated metabolic changes in breast cancer. (A) IncuCyte Zoom analysis of Δ confluency over 72 hours in a panel of breast cancer cell lines treated with NCT502 (n≥3) (B) IncuCyte Zoom analysis of Δ confluency over 72 hours of MDA-MB-468 breast cancer cells treated with NCT502 (10 µM) or PHGDH inactive control compound (10 µM) (n≥8) (C) Correlation of change in extracellular acidification rate (ECAR) (ΔECAR) in a panel of *IKBKE*-silenced breast cancer cell lines, as measured using Seahorse XF96 analysis, and the Δ confluency upon treatment of the panel of cell lines with NCT502 for 72 hours, as measured using the IncuCyte Zoom. (D) IncuCyte Zoom analysis of Δ confluency over 96 hours in a panel of breast cancer cell lines treated with DON (n≥3) (E) Correlation of change in extracellular acidification rate (ECAR) (ΔECAR) in a panel of *IKBKE*-silenced breast cancer cell lines, as measured using Seahorse XF96 analysis, and the Δ confluency upon treatment of the panel of cell lines with DON for 96 hours, as measured using the IncuCyte Zoom. (F) IncuCyte Zoom analysis of Δ confluency over 72 hours in a panel of breast cancer cell lines treated with CB839 (n≥4) Data Information: Data are n ≥ 3, mean ± SEM, *p<0.05. In (A, B, D) one-way ANOVA, in (F) two-way ANOVA with Fisher’s LSD tests were applied. In (C, E) *IKBKE-*silenced cells were transfected with a pool of 4 targeting oligos, linear regression correlation coefficients (Pearson’s R, Spearman’s rho) are shown.

